# AGE READINGS AND ASSESSMENT IN COASTAL BATOID ELASMOBRANCHS FROM SMALL-SCALE SIZE-SELECTIVE FISHERY: THE IMPORTANCE OF DATA COMPARABILITY IN MULTISPECIFIC ASSEMBLAGES

**DOI:** 10.1101/2024.03.15.585164

**Authors:** Umberto Scacco, Fabiana Zanardi, Silvio Kroha, Emanuele Mancini, Francesco Tiralongo, Giuseppe Nascetti

## Abstract

Age readings and assessment of vertebral growth increments were obtained in four batoid elasmobranch species, namely *Dasyatis pastinaca*, *Raja asterias*, *Torpedo marmorata*, and *Torpedo torpedo*. Samples were obtained opportunistically from the bycatch of a size-selective fishery, such as local small-scale trammel net fishing, in the coastal waters of the Central Tyrrhenian Sea during 2019-21. We analysed the vertebrae by a simple and rapid method preventing staining phase and histological preparation to elucidate band pairs in all species studied. Consistency of age estimates was checked by controlling for agreement on band pair counts between and within observers, and by estimating the relationships between vertebral diameter and height, and body size of the animal. Based on these data, we developed a statistical routine to obtain multiple estimates of age and growth parameters for incomplete samples due to size-selective fishing. The acceptable agreement between and within readers and the increase in vertebral size with body size demonstrated the consistency of the method. Based on the results of Von Bertalanffy and Gompertz growth models, body size was a better predictor of age than vertebral size, and Gompertz models were a better estimator of age and growth parameters than Von Bertalanffy models. The estimated parameters (k and t0, kg and cg) matched the data available for the species studied in the Mediterranean Sea. In fact, *D. pastinaca*, *T. torpedo* and *R. asterias* showed the lowest (k = 0.05-0.12), intermediate (k = 0.112-0.19) and highest (k = 0.18-0.23) growth rate, respectively. Overall, the method proved effective both in delineating band pairs in vertebrae of different species by making use of only minimal optical equipment and a single reagent, and in reliably estimating the age and growth parameters of problematic samples due to size-selective fishing. The replicability of the method will help to collect comparable demographic data in similar samples from other areas of the Mediterranean, as well as in assemblages of different species from other places.

## INTRODUCTION

Since 1980, several techniques and methods have been developed to read age on the inner vertebral surface of elasmobranch species (Cailliet et al., 1983; Cailliet et al., 1986; Cailliet, 1990; Cailliet, 2015; Hoenig and Brown, 1988; Goldman et al., 2012). The literature shows that techniques for determining age in vertebral structures of elasmobranchs are generally species-specific (Kenneth et al., 2012; Campana, 2014; Başusta et al., 2017).

Vertebral morphology is critical in choosing the appropriate method, as shape and calcification can vary widely among species (Claeson and Hilger, 2011; Smith et al., 2013; James and Natanson, 2020; Pears et al., 2020). For example, deep-water species have deep-coned, poorly calcified vertebrae, in contrast to the more calcified, less acute centra of coastal species (Gennari and Scacco, 2007; Matta et al., 2017; McMillan et al., 2017). While in the former case the use of staining techniques is almost inevitable, in the latter case direct observations of growth increments are possible, along with the use of staining or other enhancement methods (Brown and Gruber, 1988; Goldman et al., 2012; Matta et al., 2017). However, such preparations are generally species-specific and can be time-consuming and/or reagent-expensive compared with a multispecies method that avoids the staining step and uses portable stereoscopic equipment.

Estimating age and growth parameters can be difficult or impractical when estimating age in elasmobranch samples that are bycatch in a size-selective fishery (Flinn and Midway, 2021). On these occasions, samples are not complete in terms of both the number of observations and the representativeness of all expected age classes for the species under consideration (Parma and Deriso, 1990). There is a large literature on VBGE fitting in different sample situations (Flinn and Midway, 2021), as well as on the use of the increase in the size of biological structures, such as otoliths (Chih, 2009; Tyszko and Pritt, 2017) and the vertebra in species of elasmobranchs to perform aging studies (Calliet, 2015). Since age growth parameters are intrinsic characteristics of the sample (Pauly, 1980), their estimation is optimal when it can be performed simultaneously on well-sized and length-structured samples (Ogle et al., 2017). Unfortunately, samples are generally far from optimal as they usually suffer from poorly represented size ranges and/or errors in age estimation (King et al., 2017), such as samples from size-selective fisheries.

The small-scale fishery is one of the most size/species selective fisheries, as it relies on the use of several highly selective passive fishing gears, such as fixed and drift nets, hooks, and pots (FAO 1972, 1985, 1995, 2016; JRC, 2020). This type of fishing is concentrated in coastal waters (Smith and Basurto, 2019), sometimes very close to shore, as in the study area, as well as in other areas of the Italian and Mediterranean seas (Tiralongo et al., 2018a; Tiralongo et al., 2020; Silva et al., 2021).

The multispecific method we developed was tested on four batoid elasmobranch coastal species typical of soft sandy-muddy bottoms of the Mediterranean Sea (Bauchout 1997). *Dasyatis pastinaca* (Linnaeus, 1758), *Raja asterias* Delaroche, 1809, *Torpedo marmorata* Risso, 1810 and *Torpedo torpedo* (Linnaeus, 1758), are emblematic of the bycatch of different metiers in the Mediterranean Sea, such as bottom trawl (Barria et al., 2015) and small-scale fishery (Tiralongo et al., 2018a; Lloret et al., 2019). These species live in inshore and brackish waters, such as the common sting ray *D. pastinaca* (Brito, 1991; Seret, 2003). The common stingray is an opportunistic feeder (Tiralongo et al., 2020) and an aplacental viviparous species (Dulvy and Reynolds, 1997). In contrast, the starry ray *R. asterias* is an oviparous (Dulvy and Reynolds, 1997) and coastal endemic species to the Mediterranean, particularly the western sector (Fatimetou and Younes, 2016). The starry ray is an opportunistic feeder species (Stehmann and Bürkel, 1984) and has some commercial importance, albeit low, in the central Tyrrhenian Sea, particularly in the landings of the trawl fishery (Romanelli et al., 2007). The common torpedo *T. torpedo* and the electric marbled ray *T. marmorata* are sympatric species with slightly different bathymetric distribution and biological characteristics (Consalvo et al., 2007). The common torpedo inhabits shallower depths than the electric marbled ray (Capapé and Desoutter, 1990), but both species share a piscivorous feeding strategy (Møller, 1995; Barria et al., 2015; Tiralongo et al., 2019) and aplacental viviparity (Mellinger, 1971; Last et al., 2016). The electric marbled ray inhabits a more heterogeneous habitat (Michael, 1993) and grows larger than the common torpedo, with a common length of about 60 cm (Last et al., 2016) and up to 100 cm maximum total length recorded (Mller, 1995).

The aim of this work is to provide age and growth estimates in a multi-specific assemblages of coastal batoid elasmobranchs by a unique and rapid method for reading and assessing age in problematic samples that are usual bycatch of size-selective coastal fisheries. It can help obtain reliable and comparable age and growth data for coastal elasmobranch species.

## MATERIAL AND METHODS

### Sample collection

All samples were collected through fishery-dependent surveys (on seven occasions between October 2019 and March 2021) planned at the Montalto di Castro (Central Tyrrhenian Sea, Italy) small-scale fisheries landing site in the Fiora harbor-canal (Figure 1). The species sampled were from the bycatch of trammel nets used in coastal waters in the area (Figure 1) in a depth range of 5 to 20 m. The net was about 2 km long, with a mesh size of 3 cm for the inner panel and 10-15 cm for the outer panel. The trammel net was usually lowered at sunset and hauled at dawn. We trained fishermen to facilitate the release of live specimens into the sea whenever possible and safe, and to collect all dead elasmobranch samples despite their size. After collection, specimens were immediately taken to the laboratory and stored at −20°C until dissection.

**Fig. 1.**
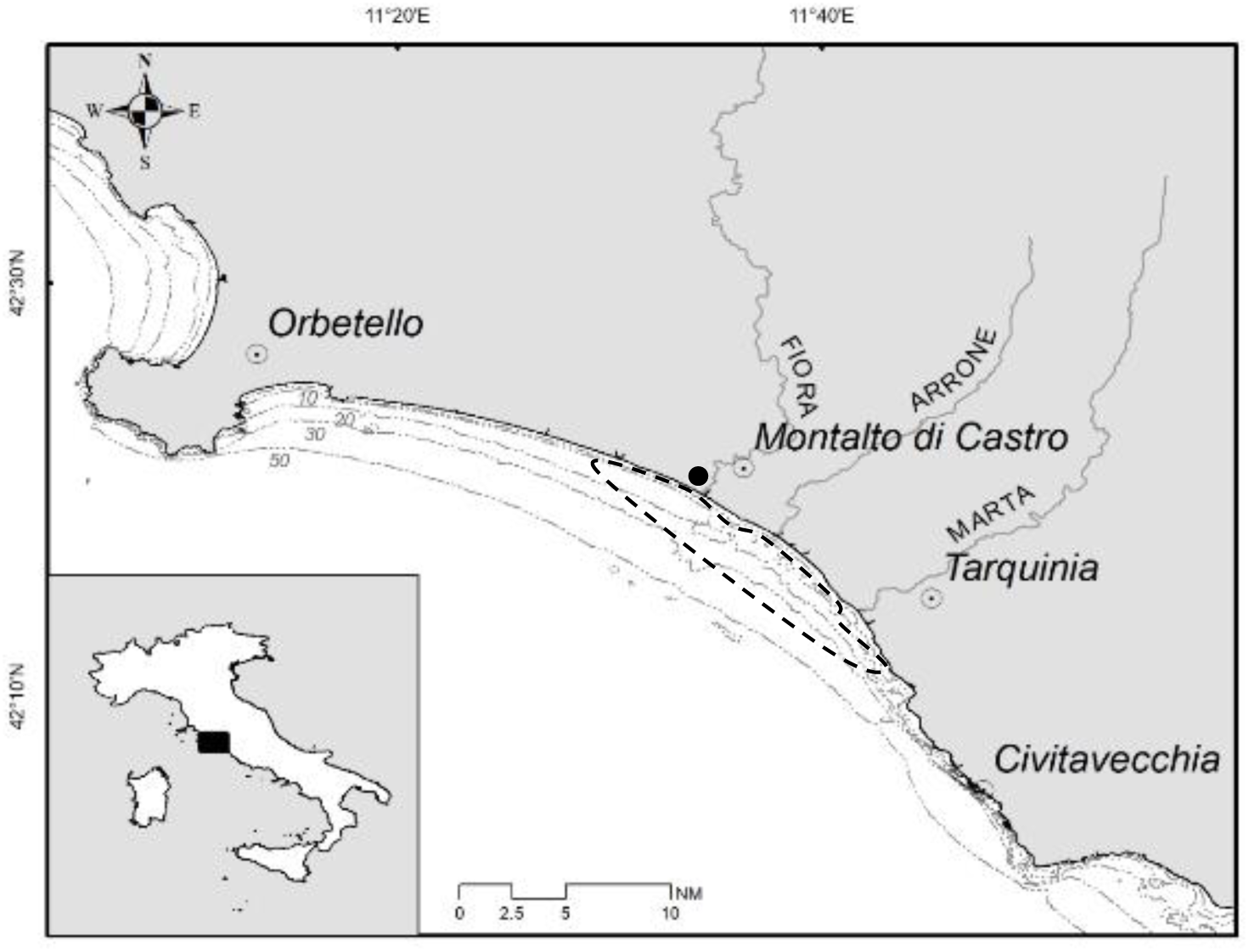
Map of the study area (central Tyrrhenian Sea, Italy, Mediterranean) with bathymetry, principal river mouths and towns. Dotted figure encloses the area where samples came from and dark circle indicates local landing site chosen for sampling.

### Biometrics

Specimens were identified taxonomically according to Bauchot et al. (1987) and Serena (2005) and then measured and weighed to the nearest mm and mg for total length (TL) and disc width (DW) and weight, respectively. TL was not recorded for *D. pastinaca* because of the often-damaged tails. The specimens were sexed and the masses of liver, stomach, uterus and dissected ovaries were weighed to the nearest mg. Sexual maturity was assessed on the scale developed by Follesa and Carbonara (2019). A digital scale and a metal ichthyometer (scale sensitivity 1 mg and 1 mm, respectively) were used to collect biometric data.

### Laboratory preparation of vertebrae

We removed a postcranial portion of the vertebral column approximately 5 cm long, cutting away excess organic tissue as best we could. Based on a combined approach between vertebral preparations proposed by Hoenig and Brown (1988) and Gennari and Scacco (2007), we performed several trials to test the effectiveness of the method prior to reading and age assessment.

Specifically, samples were tested to determine the bleach soak time necessary to remove organic tissue residues without compromising the growth increment reading. Once the optimal protocol was established, we followed the following steps:

1. The column section was immersed in NaClO (bleach) solution for between 30 and 60 minutes (larger sections took longer than smaller ones).
2. The column section was then rinsed with distilled water and sectioned into individual vertebrae.
3. The individual vertebrae were re-immersed in a new NaClO solution for 5-10 minutes depending on size, shaking them slowly to allow the solution to penetrate and remove residual tissue. Too short a time does not effectively remove all connective tissue, while too long a time literally destroys the vertebra.
4. Once removed from the solution, the vertebrae were rinsed with distilled water and allowed to dry under the laboratory fume hood.
5. For age determination, each vertebra was cut sagittally with a sharp scalpel. In this way, we obtained two halves (two half hourglasses for short) in which to observe the inner concave surface in which the periodic increments are arranged in series, from the vertebral centre toward the margin. Next, the two halves were placed on a plate and immersed in simple glycerol (C3H8O3) to obtain synoptic observation of the symmetrical sides simultaneously. Next, the samples were observed with the Dino-Lite digital microscope (online information 1 a, b), in a dark room. The above transmitted LED light allowed us to observe the different contrast growth increments best visualized when hit by the multi-beam incident light oriented orthogonally of the sample. We added an external warm light lamp used laterally to enhance the contrast of the bands.
6. Measurements of vertebral diameter (VD) and height (VH) of individuals were made using Dino-Lite digital microscope software on images transmitted directly to a personal computer with an accuracy of 0.001 mm (Figure 2).
7. Two readers independently performed age counts twice each on images selected in random order and without reference to specimen size for each species considered. We counted one opaque and one translucent band as a periodic increment. We considered a distinct band for counting when it was distinguishable from adjacent bands and observed continuously and correspondingly between the two sides of the sagittal section of the vertebra, on its inner surface.

**Fig. 2.**
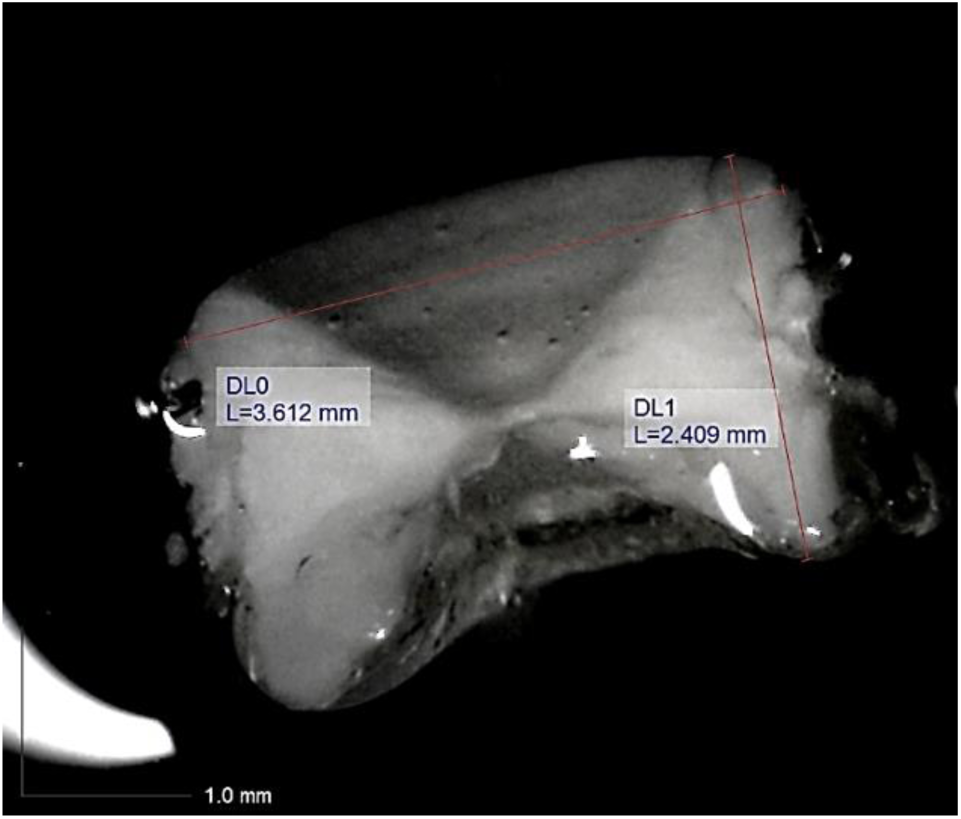
Example of the measures (red perpendicular lines, L= size in mm; DL0= vertebral diameter; DL1= vertebral height) taken on a vertebral image obtained through observation under Dino-Lite equipment of an individual of *Dasyatis pastinaca* measuring 380 and 230 mm in total length and disk width, respectively (snapshot 1, acquisition date 15/06/2020)

### Statistical methods

We checked intra- and inter-reader agreement on age estimates by the average percent error index (IAPE, Beamish and Fournier, 1981) on each species separately. We regressed VD and VH, as well as their ratio (VD/VH), on DW and TL using a linear model of the type

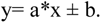

A test of parallelism was used in pairwise comparisons of the slopes of the regressions between species for each variable and predictor, separately. In the pairwise multiple comparisons, the p-level for acceptance of the results was lowered by Bonferroni’s correction (Zar, 1999). The normality of the size frequency distributions of the studied species was checked by Liliefors, Komolgorov-Smirnov and Shapiro’s tests. The difference between the sex ratios of the species was checked through separate chi-square tests. Because of the incomplete range of sample sizes (consisting mostly of juveniles and subadults), we performed separate bootstrapping procedures of random resampling with replacement of individual length data (such as VH, VD, DW, and TL) according to age in the original samples of each species. The number of iterations was set at 1000 to ensure a stabilized cumulative mean as close to the true sample mean as possible. We then used the analytic geometric method of Ford (1933)-Walford (1946) replicated on each generated sample to obtain the overall mean asymptotic lengths provided with error (std. err, 95% conf. limits) over 1000 iterations. In this task, we used the models with 0 < slope < 1 and intercept > 0. In fact, negative slopes equal to and greater than 1 imply a lack of fit to estimate asymptotic dimensions (Ford, 1933; Walford, 1946); as do distant models with intercept ≤ 0. Finally, we used these values as fixed asymptotic parameters in the Von Bertalanffy logistic equations (Von Bertalanffy L 1938).

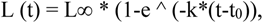

and in the Gompertz model (Gompertz, 1825)

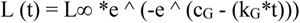

to fit the corresponding length-at age data and calculate the parameters k, t_0_, k_G_ and c_G_ with their error intervals, where

k is the rate of growth in length with increasing age,

t_0_ is the estimated time at birth,

k_G_ is the Gomperz rate of growth, and

c_G_ is the Gomperz parameter.

The estimation method and loss function were the Levenberg-Marquardt algorithm and least squares minimization for linear and nonlinear fit, respectively. Due to the limited sample size, unbalanced sex ratio and uneven age distribution, growth parameters were not estimated for *T. marmorata*. Statistical analyses were performed with STATISTICA 7.1, and Office Excel version 18.2110.13110.0 was used to perform bootstrapping on the original samples by resampling with replacement through the command:

INDEX($first sample$first row:$last sample$last row;ROWS($first sample$first row:$last sample$last row)*RANDOM()+1).

## RESULTS

The agreement between the two independent readings showed an error range that can be considered acceptable (< 5%) for all studied species, both inter- and intra-reader (Table 1). However, the IAPE was different among the species (Table 1). *T. torpedo* showed the lowest value, *R. asterias* and *T. marmorata* the intermediate value and *D. pastinaca* the highest IAPE (Table 1). The distribution of the error also differed as the age of the species considered increased. The error occurred randomly in 12 specimens of *D. pastinaca* of different sizes, with a difference of 1 year between readers in all cases except 2 years in the largest specimen (Figure 3 a). *Raja asterias* showed a similar pattern (7 individuals), with a difference of 1 year in all cases (Figure 3 b). In *T. torpedo*, the error was concentrated in the large individuals, but occurred in a few of them (3), with a difference of 1 year (Figure 3 d). Similarly, the error increased with size in *T. marmorata*, with a 2-year difference between readers in the largest specimen (Figure 3 c).

**Figure 3.**
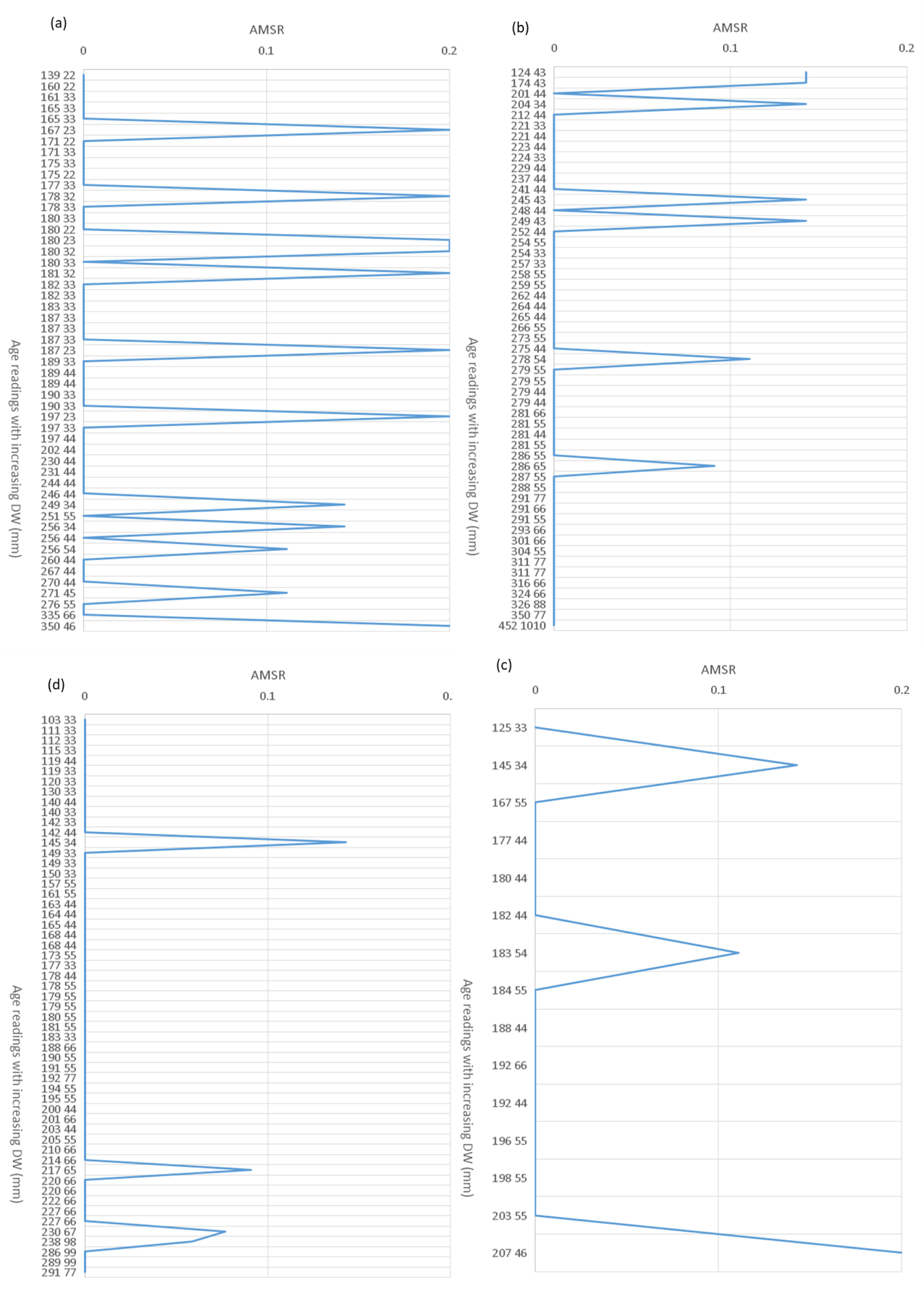
**a, b, c, d** Age-bias plots representing the variation of the Absolute values of Mean-Standardized Residuals (AMSR) between age readings (as couples of numbers placed on the right of the size in cm) of two independent readers with increasing body size as disk width (DW, mm) of four coastal batoid species sampled in the central Tyrrhenian Sea: (a) *Dasyatis pastinaca*, (b) *Raja asterias*, (c) *Torpedo marmorata* and (d) *Torpedo torpedo*.

**Table 1.** Index of Average Percent Error (IAPE) calculated between and within two readers based on two independent age readings within each reader. Data refer to a fishery-dependent sample of four coastal batoid species assemblage from the central Tyrrhenian Sea.

| IAPE by Species | Intra-reader 1 | Intra-reader 2 | Inter-readers |
| --- | --- | --- | --- |
| *Dasyatis pastinaca* | 3.33% | 2.91% | 4.26% |
| *Raja asterias* | 0.92% | 1.02% | 1.67% |
| *Torpedo marmorata* | 1.48% | 2.08% | 2.81% |
| *Torpedo torpedo* | 0.16% | 0.59% | 0.73% |

Photographic images showed that the contrast and sharpness of the bands varied among species. Common stingrays had weaker bands (Figs. 4 a, b, c), particularly in older specimens (5-6 years old) (Fig. 4 c), compared to the other species. *D. pastinaca* had poorly contrasted outer bands, with thin opaque areas and large intermediate areas that were poorly translucent. In the case of the starry skate (Figs. 4 d, e, f), the opaque and translucent zones had greater contrast than in the earlier species. However, in the oldest individual (Fig. 4 f) of this species, some degree of folding of the outer margin was noted. Periodic increments were clear in the common torpedo, with good contrast and sharpness (Figs. 4 g, h, i). A similar situation was observed for the marbled electric ray (Figs. 4 j, k, l), with darker opaque bands in young and intermediate specimens (Figs. 4 j, k) than in older ones. Birthmark was observed particularly in the sectioned vertebrae of *T. marmorata* and *R. asterias*, as a change in angle of the corpus calcareum was noted in small (Fig. 4 d and j, respectively), intermediate (Fig. 4 e and k, respectively) and large individuals (Fig. 4 f and l, respectively) of these species.

**Fig. 4.**
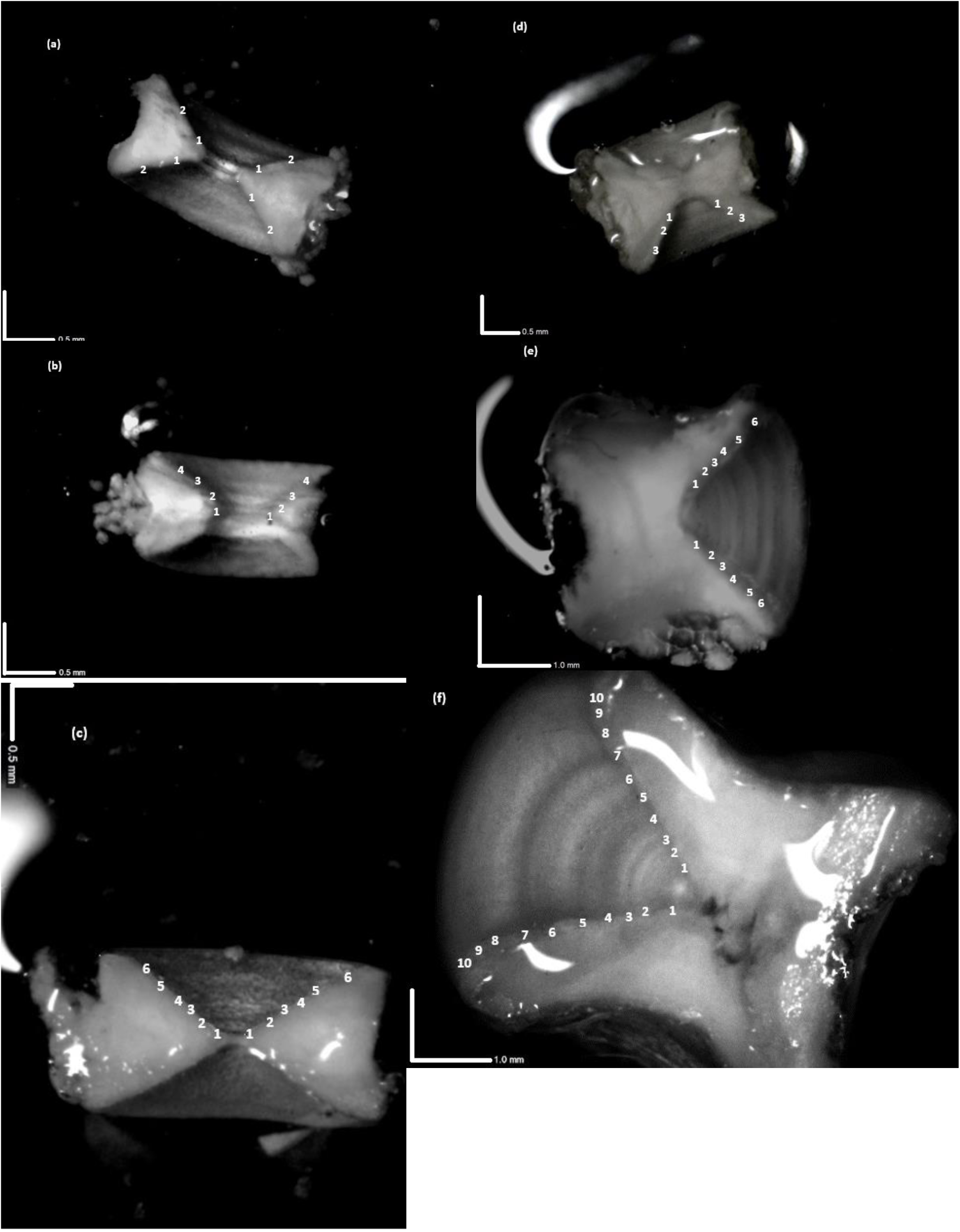
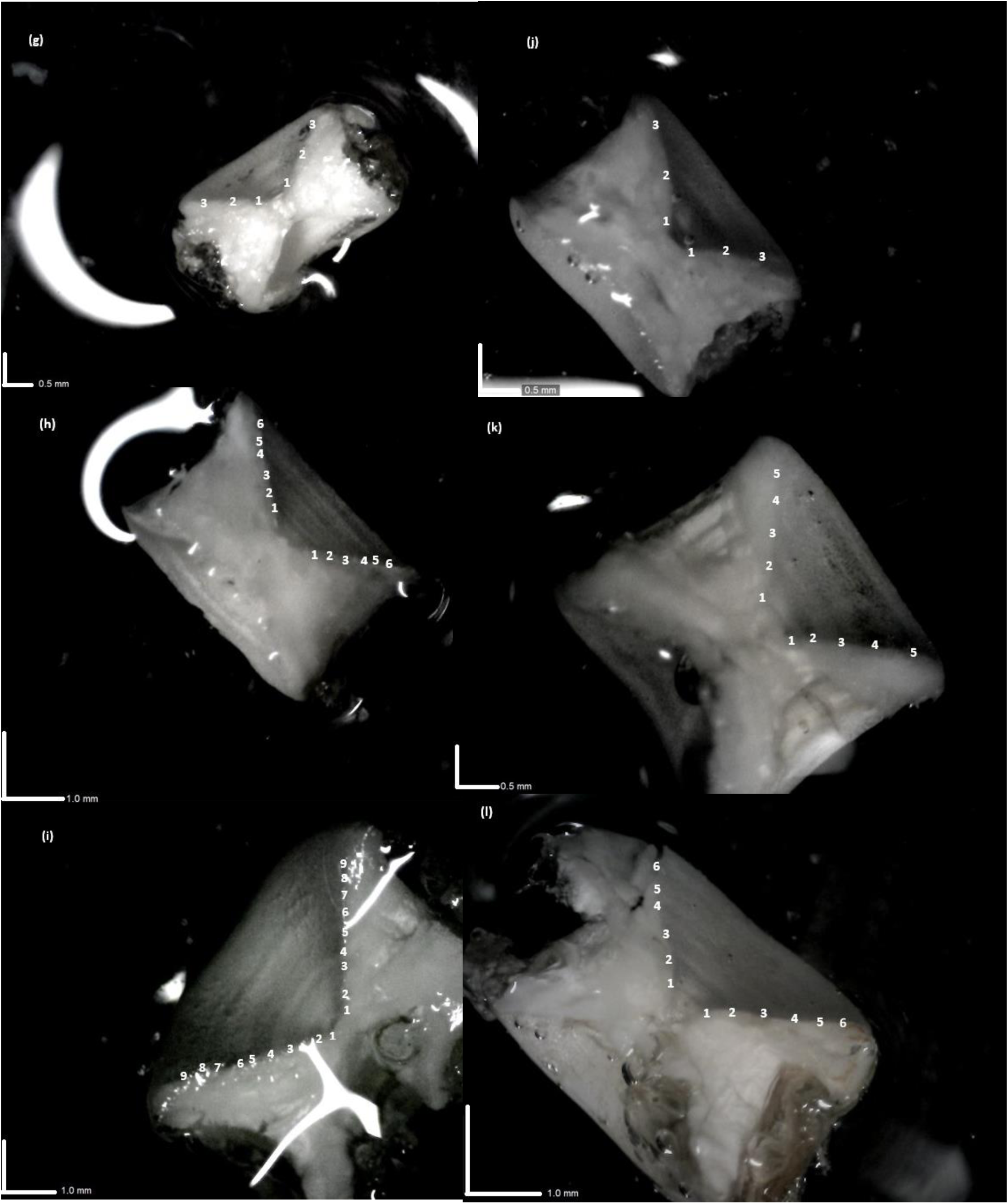
**a, b, c, d, e, f, g, h, I, j, k, l** Sagittal sections of vertebrae of four coastal batoid elasmobranch species observed in glycerol immersion under incident led light by Dino-Lite equipment with indication of rings and corresponding age counts. *Dasyatis pastinaca* (a, b, c), *Raja asterias* (d, e, f), *Torpedo torpedo* (g, h, i), *Torpedo marmorata* (j, k, l). Bold white scales bars are reported. Juveniles (a, d, g, j), intermediate (b, e, h, k) and large (c, f, i, l) individuals are shown. Magnification: a = 79.4 x, b = 81.7 x, c = 62.3 x, d = 59.9 x, e = 55.4 x, f = 55.4 x, g = 61.5 x, h = 43.5 x, i = 40.2 x, j = 62.3 x, k = 58.6 x, l = 51.9 x

The goodness of fit of the linear model varied across species and variables (Table 2, Online Information 2). However, the relationships showed a general linear increase in diameter and vertebral height with increasing DW (Figs. 5 a, b, c, d) and TL (Online Information 3 a, b, c) in all species, although not significant in *T. marmorata* (Table 2, Online Information 2). The common torpedo showed the steepest slopes for both VD and VH along DW and TL (Fig. 5 c, Online Information 3 c; Table 2, Online Information 2). In the other cases, regression slopes varied according to species, vertebral size, and predictor. For example, *R. asterias* (Online Information 3 a) and *D. pastinaca* (Fig. 5 a) showed the lowest slopes for the VD-TL and VH-DW regressions, respectively (Table 2, Online Information 2). The VD/VH ratio followed an invariant trend with increasing size, or slightly decreasing in some species (Figs. 5 a, b, c, d; online information 3 a, b, c). For this ratio, we observed very low explained variances and never significant slopes in all species except *D. pastinaca* (Fig. 5 a, Table 2, online information 2).

**Fig. 5.**
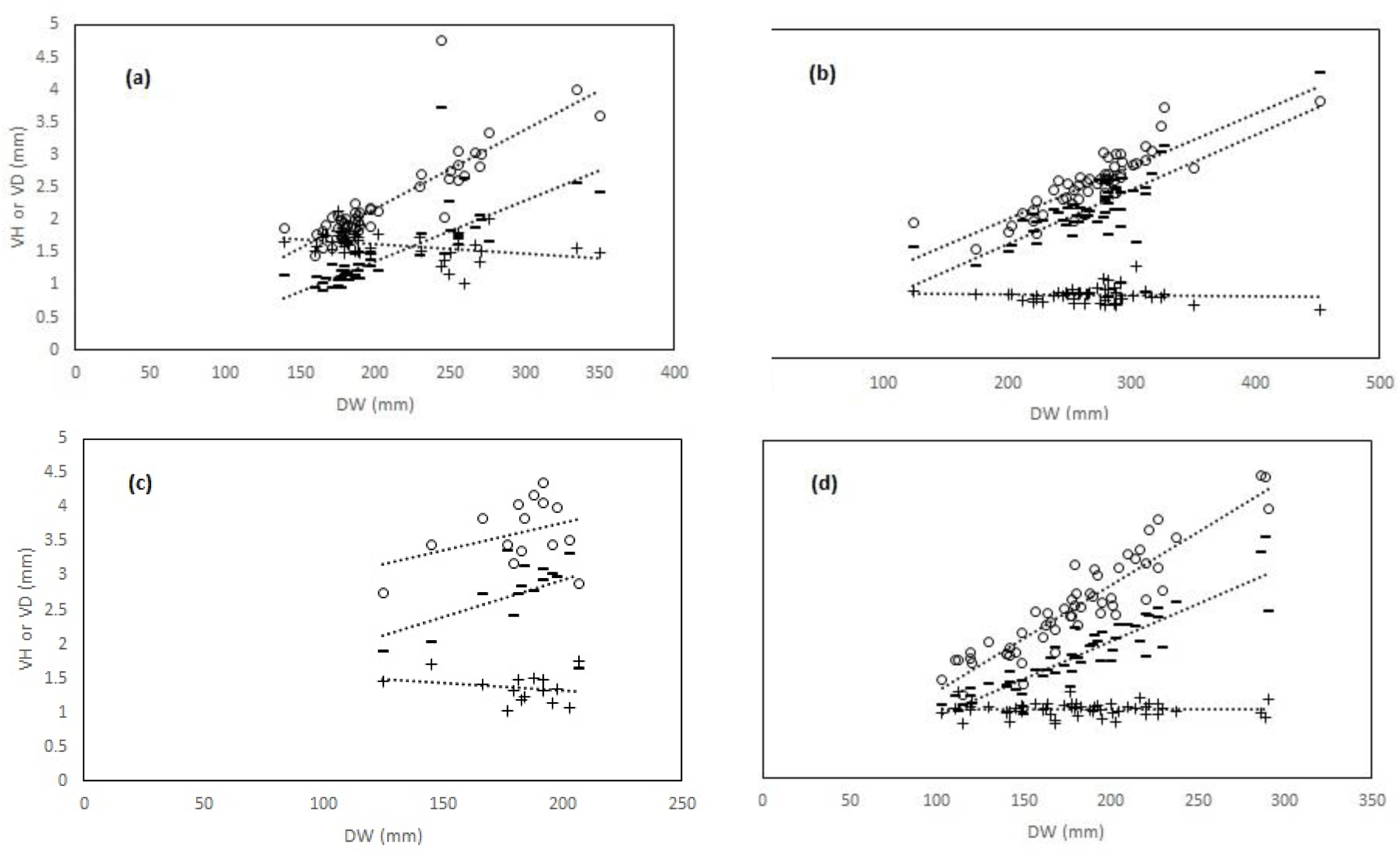
**a, b, c, d** Linear relationships (dotted lines) between disk width (DW) and vertebral measures (VD vertebral diameter, empty circles; VH vertebral height, dark lines; VD/VH ratio, dark crosses) in four coastal batoid elasmobranch species form the central Tyrrhenian Sea. (a): *Dasyatis pastinaca*; (b): *Raja asterias*; (c): *Torpedo marmorata*; (d): *Torpedo torpedo*

**Table 2.**
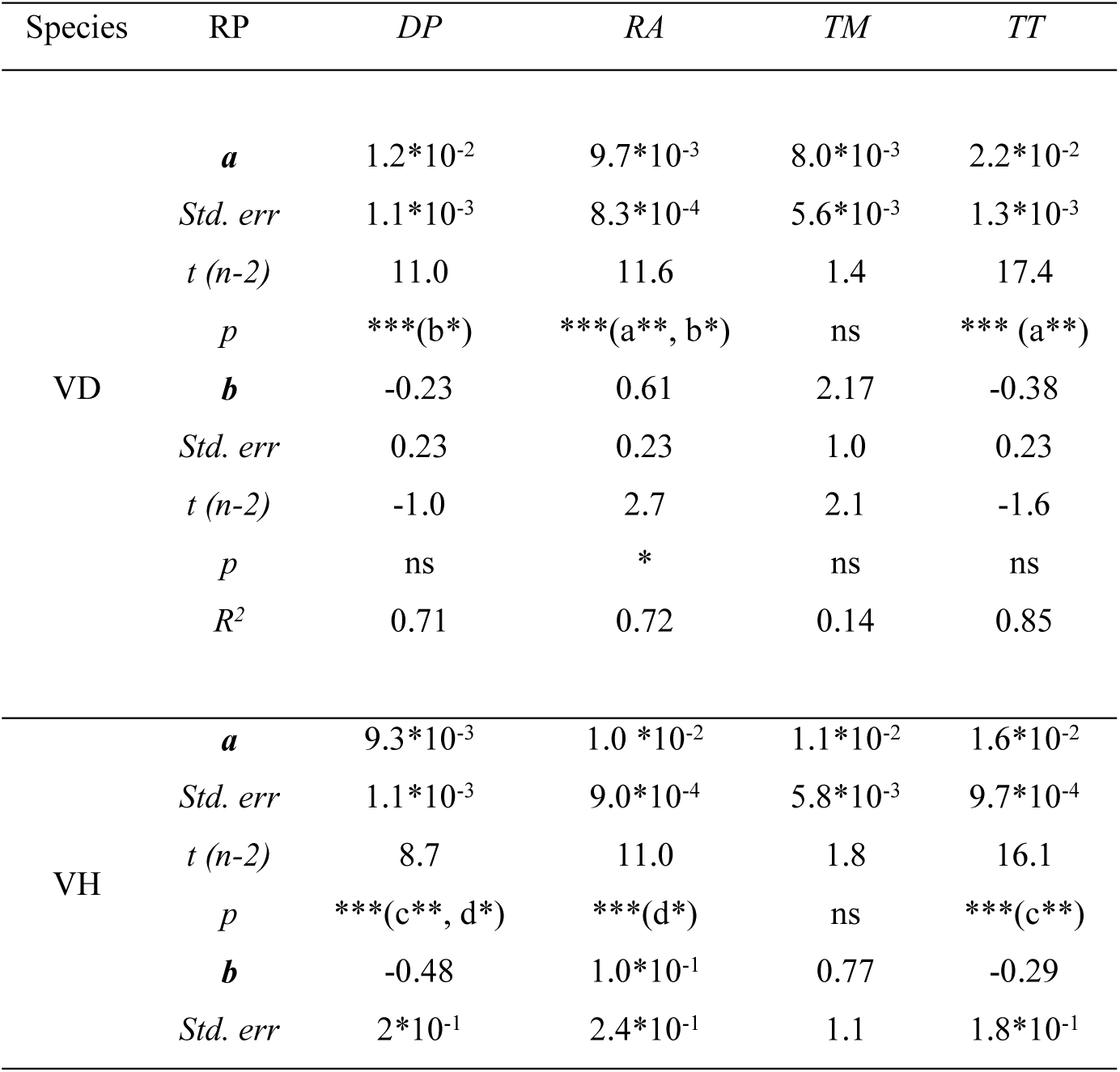
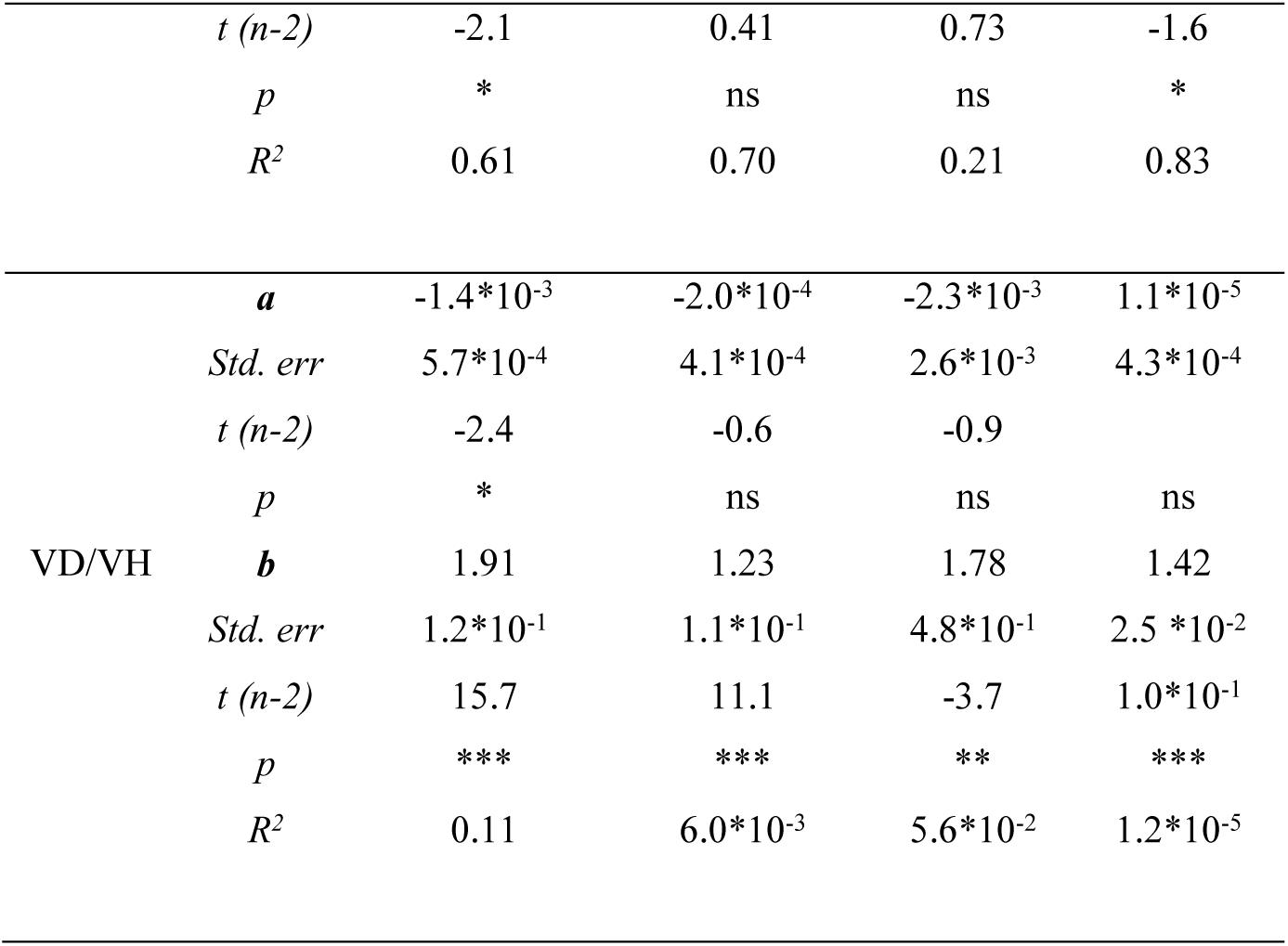
Regression parameters (RP, a: slope, b: intercept, R^2^: explained variance, *Std. err* : standard error; *t (n-2)*: degree of freedom of t statistic; and p: level of significance) of linear relationships between vertebral dimensions and increasing fish size as disk width (mm) of four coastal batoid species (DP: *Dasyatis pastinaca*, RA: *Raja asterias*, TM: *Torpedo marmorata* and TT: *Torpedo torpedo*) sampled in the central Tyrrhenian Sea. VD, VH and VD/VH are vertebral diameter, height and their ratio, respectively. Same letter in brackets indicates significantly different regression slope between species with asterisks denoting its p-level of significance. ns stands for not significant differences. * p < 0.05, ** p < 0.01, *** p < 0.001

The size frequency distributions (Online information 4 a, b, c, d) of the studied species had different deviances from normality. In the case of *D. pastinaca* (Online information 4 a), distribution was right-skewed (asymmetry: 1.30 ± 0.33 std. err.) with respect to the sample mean DW (206.78 mm ± 6.32 std. err.) and not normal (K-S: d=0.25, p<0.01; Lilliefors, p<0.01; W Shapiro-Wilkins, W=0.84, p<0.001). Three different cohorts may be argued in this species, with observations more concentrated in the smaller compared to larger size class (Online information 4 a). Differently, the distributions for *R. asterias* (K-S, d =0.13, p > 0.05; Lilliefors, p <0.05; Shapiro-Wilk, W = 0.92, p <0.01) and *T. marmorata* (K-S, d= 0.22, p > 0.05; Lilliefors, p < 0.05; Shapiro-Wilk, W = 0.86, p < 0.05) appeared closer to normality compared to *D. pastinaca*. In fact, we found a moderate right (0.46 ± 0.33 std. err.) and left skewed asymmetry (−1.53 ± 0.58) with respect to the corresponding sample means (268.26 mm ± 6.48 std. err. and 181.27 mm ± 5.61 std. err.) in the distributions of *R. asterias* and *T. marmorata*, respectively. Therefore, observations were more concentrated in the larger and smaller size classes in *T. marmorata* and *R. asterias*, respectively (Online information 4 b, c). The distribution was normal in the case of *T. torpedo* (K-S d=0.05, p > 0.05; Lilliefors, p > 0.05; Shapiro-Wilk, W = 0.96, p > 0.05), with negligible right skewness (asymmetry 0.32 ± 0.41 std. err.) with respect to the mean of the sample (179.82 mm ± 5.86 std. err.). In this case, the observations appeared to be roughly equally distributed around the mean (Online information 4 d). The sex ratios were balanced in *D. pastinaca* (F/M = 0.83, X^2^ = 0.49, d.f. = 1, p > 0.05) and *R. asterias* (F/M = 0.96, X^2^ = 0.02, df = 1, p > 0.05), while males were exclusive in *T. marmorata* and females prevailed on males in *T. torpedo* (F/M = 2.05, X^2^ = 6.56, d.f. = 1, p < 0.01).

Out of a total of 1000 iterations performed for all species and variables, the cumulative sample averages reached stability after a different number of cumulative iterations, depending on the species and variable involved (online information 5 and 5 bis, a, b, c, d). For example, the number of iterations needed to stabilize the sample mean of *T. torpedo* was generally higher for all variables than for the other species (online information 5 and 5bis, c). In contrast, the cumulative sample averages of VD and VH in *T. marmorata* remained unstable around and beyond the maximum number of iterations performed, respectively (online information 5 d).

The number of appropriate Ford-Walford fits (0 < slope < 1 and intercept > 0) to compute the mean asymptotic lengths differed depending on the species and variable in question, but with no apparent ordering along variables and/or species (Table 3). In terms of the error of the mean asymptotic estimates, it was acceptably narrow in all species for all variables, but in *D. pastinaca* for VH ∞ and DW∞. *Torpedo marmorata* showed lower errors along all variables than the other species (Table 4). Nevertheless, the estimates of asymptotic lengths were not realistic for this species (Table 3). Therefore, we excluded *T. marmorata* from further analysis.

**Table 3.**
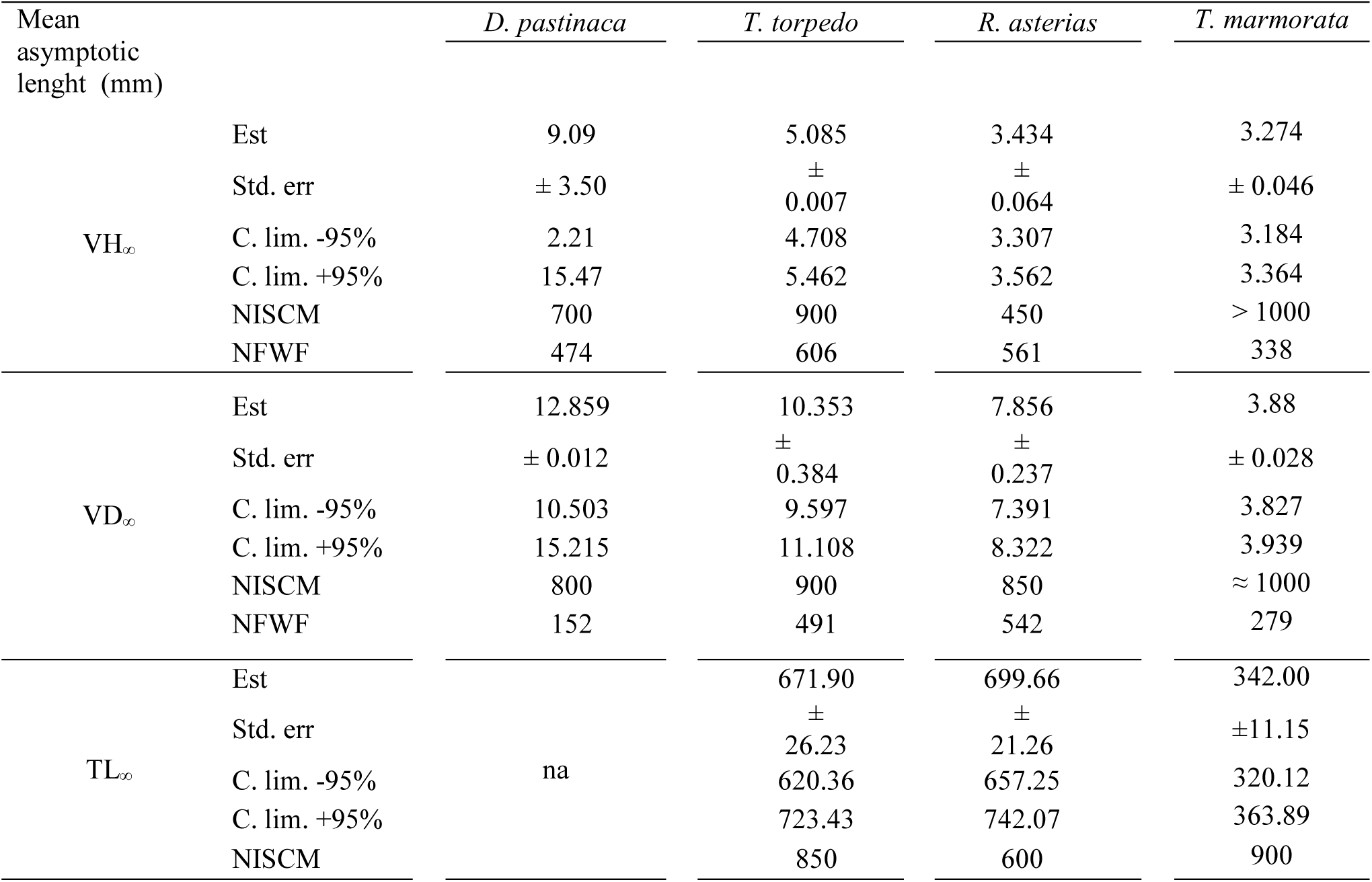
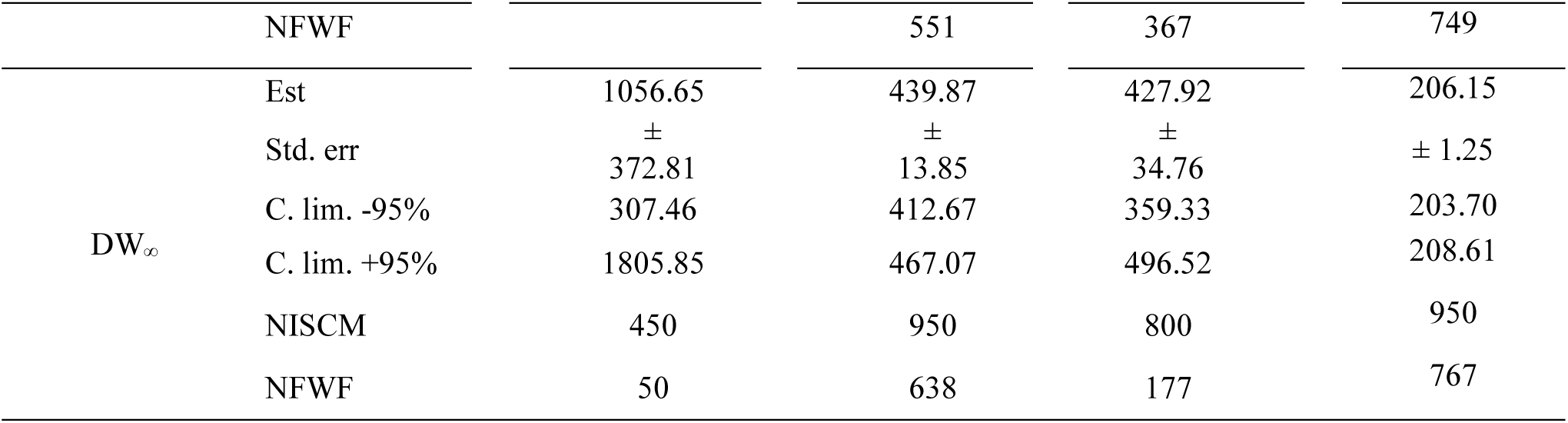
Errors (standard error and 95% upper and lower Confidence Limits) of mean asymptotic lengths expressed as vertebral height (VH), diameter (VD), total body length (TL) and disk width (DW) and estimated through Ford-Walford fits over 1000 bootstrapping iterations (NISCM: approximate Number of Iterations needed to Stabilize Cumulated Mean length; NFWF: Number of Ford-Walford Fits used to calculate mean asymptotic length) on original data of four coastal batoids elasmobranchs from the central Tyrrhenian sea

**Table 4.** Errors (±Std. err.), estimated growth parameters (k and t_0_: VBGE rate of growth and time at size 0, respectively; k_g_ and c_g_: Gompertz rate of growth and parameter, respectively) and statistics (R^2^ and t _n-2_: explained variance and t statistic for n-2 degrees of freedom with n as sample size, respectively) of VBGE and Gompertz models calculated by imposing mean bootstrapped asymptotic lengths (values right to numbers in superscript brackets indicating correspondence with species) as vertebral height (VH∞), diameter (VD∞), total length (TL_∞_) and disk width (DW_∞_) for three species of coastal batoids elasmobranchs from the central Tyrrhenian sea. ** p< 0.01, *** p< 0.001, borderline p-levels are in brackets. Ns: not significant p > 0.05. Numbers in normal brackets indicate sample size by species.

|  |  | <sup>(1)</sup> *D. pastinaca* (53) |  |  |  | <sup>(2)</sup> *T. torpedo* (55) |  |  |  | <sup>(3)</sup> *R. asterias* (51) |  |  |  |
| --- | --- | --- | --- | --- | --- | --- | --- | --- | --- | --- | --- | --- | --- |
|  |  | VBGE |  | GOMPertz |  | VBGE |  | GOMPertz |  | VBGE |  | GOMPertz |  |
| | | k | $t_0$ | $k_g$ | $c_g$ | k | $t_0$ | $k_g$ | $c_g$ | k | $t_0$ | $k_g$ | $c_g$ |
| $VH_\infty$<br>(1) 9.09<br>(2) 5.08<br>(3) 3.43 | Est | 0.049 | -0.153 | 0.138 | -1.081 | 0.182 | 0.774 | 0.267 | -0.882 | 0.378 | 0.201 | 0.432 | -0.441 |
| | ±Std. err. | 0.008 | 0.605 | 0.084 | 0.023 | 0.016 | 0.305 | 0.022 | 0.102 | 0.093 | 0.926 | 0.106 | 0.431 |
| | $t_{n-2}$ | -5.85 | 0.25 | 12.92 | 6.05 | 11.46 | 2.54 | 11.84 | 8.61 | 4.06 | 0.22 | 4.07 | 1.02 |
|  | p | *** | ns | *** | *** | *** | * | *** | *** | *** | ns | *** | ns |
| | $R^2$ | 0.42 | | 0.42 | | 0.77 | | 0.78 | | 0.40 | | 0.41 | |
| $VD_\infty$<br>(1) 12.86<br>(2) 10.35<br>(3) 7.86 | Est | 0.047 | -0.767 | 0.127 | -0.983 | 0.093 | 0.091 | 0.163 | -0.835 | 0.064 | -3.491 | 0.102 | -0.375 |
|  | Std.d err | 0.007 | 0.586 | 0.017 | 0.061 | 0.008 | 0.363 | 0.013 | 0.064 | 0.008 | 1.103 | 0.013 | 0.065 |
| | $t_{n-2}$ | 7.05 | -1.30 | 16.13 | 7.56 | 12.24 | 0.25 | 12.60 | 13.10 | 7.36 | -3.16 | 7.64 | 5.76 |
|  | p | *** | ns | *** | *** | *** | ns | *** | *** | *** | ** | *** | *** |
| | $R^2$ | 0.51 | | 0.53 | | 0.76 | | 0.77 | | 0.54 | | 0.55 | |
| $TL_\infty$<br>(2) 671.9<br>(3) 699.7 | Est | | | | | 0.113 | -0.264 | 0.177 | -0.679 | 0.125 | -2.188 | 0.168 | -0.193 |
|  | Std.d err |  |  |  |  | 0.009 | 0.391 | 0.014 | 0.069 | 0.015 | 0.809 | 0.020 | 0.092 |
| | $t_{n-2}$ | | | na | | 12.05 | -0.67 | 12.27 | 9.90 | 8.23 | -2.70 | 8.49 | 2.11 |
|  | p |  |  |  |  | *** | ns | *** | *** | *** | ** | *** | * |
| | $R^2$ | | | | | 0.76 | | 0.77 | | 0.63 | | 0.64 | |
| $DW_\infty$<br>(1) 1056.6<br>(2) 439<br>(3) 427.9 | Est | 0.048 | -1.227 | 0.121 | -0.899 | 0.102 | -0.574 | 0.162 | -0.643 | 0.173 | -1.088 | 0.222 | -0.269 |
|  | Std.d err | 0.004 | 0.442 | 0.010 | 0.039 | 0.009 | 0.444 | 0.014 | 0.067 | 0.025 | 0.777 | 0.031 | 0.141 |
| | $t_{n-2}$ | 10.36 | .2.77 | 11.46 | 22.63 | 11.40 | -1.29 | 11.60 | 9.62 | 7.00 | -1.40 | 7.12 | 1.91 |
|  | p | *** | ** | *** | *** | *** | ns | *** | *** | *** | ns | *** | (0.06) |
| | $R^2$ | 0.70 | | 0.72 | | 0.74 | | 0.74 | | 0.57 | | 0.58 | |

On the basis of the results of both VBGE and Gompertz models, the mean explained variance of fits was generally lower when vertebral lengths were used as fixed asymptotic parameters rather than body lengths (Table 4). This was observed in all species except *T. torpedo*, which showed high R^2^ despite the model and variable used (Table 4). In general, Gompertz produced better (negative) estimates of t_0_ than the VBGE models for all species and variables (Figs. 6, 7 a, b, c, d; Table 4). In fact, VBGE models produced positive t_0_ estimates when vertebral measures VH and VD were used as asymptotic lengths in the models of *R. asterias* and *T. torpedo* (Fig. 6 a), and *T. torpedo* (Fig. 6 b), respectively (Table 4). The growth rates of species (kg) were higher in the Gompertz compared to the VBGE models (k) across all variables (Figs. 6, 7 a, b, c, d; Table 4). In general, the range of growth rates estimated had the highest values in *R. asterias* (k and kg = 0.18 - 0.23), the intermediate ones in *T. torpedo* (0.12 - 0.19), and the lowest in *D. pastinaca* (0.05 - 0.12), comparing all variables and models across species (Figs. 6 a, c, d, 7 a, b, c, d; Table 4).

**Fig. 6.**
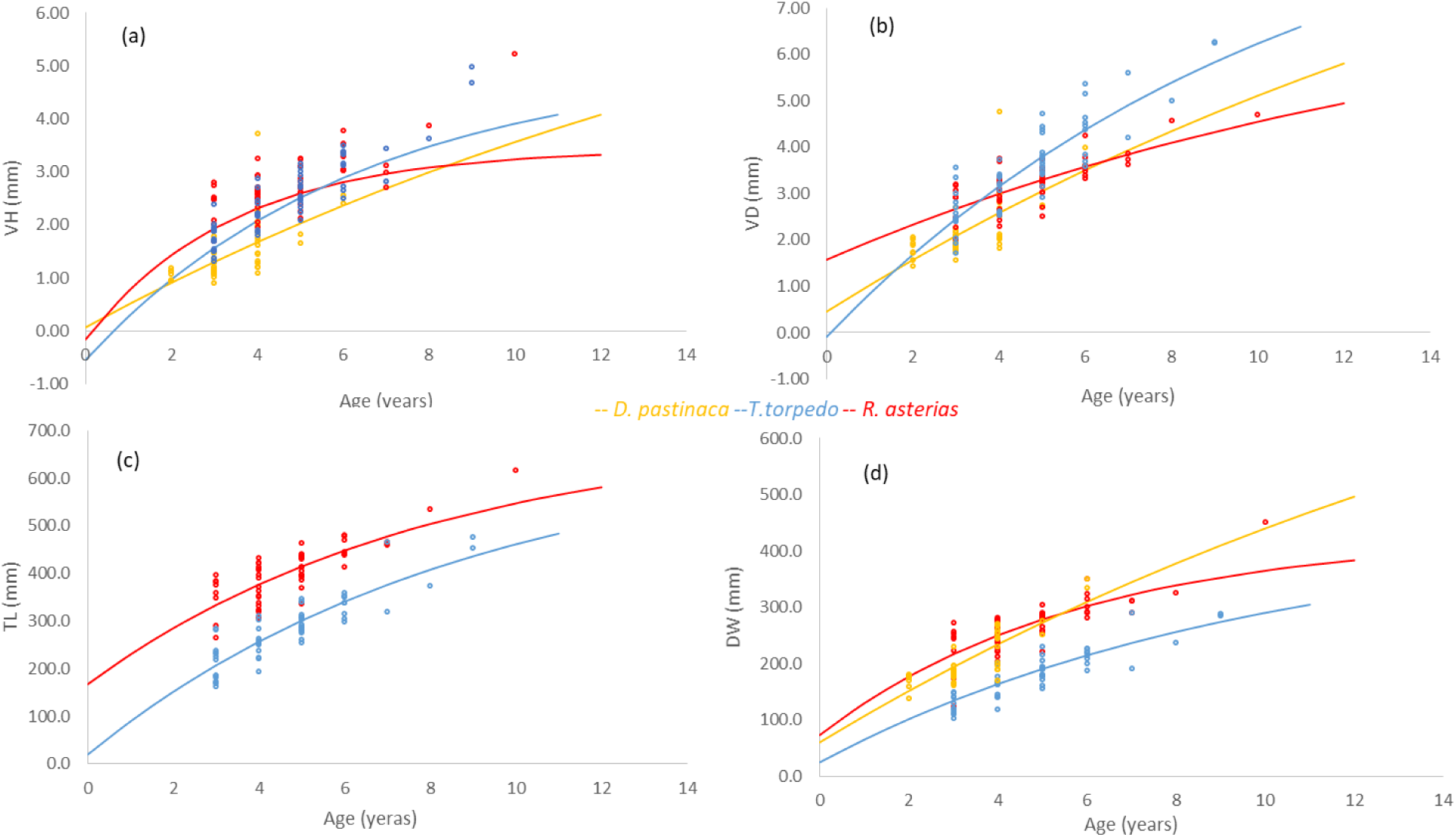
**a, b, c, d** VBGE models based on vertebral (a) height (VH) and (b) diameter (VD), (c) total length (TL) and (d) disk width (DW) used as asymptotic lengths in *Dasyatis pastinaca*, *Raja asterias* and *Torpedo torpedo* from the central Tyrrhenian Sea

**Fig. 7.**
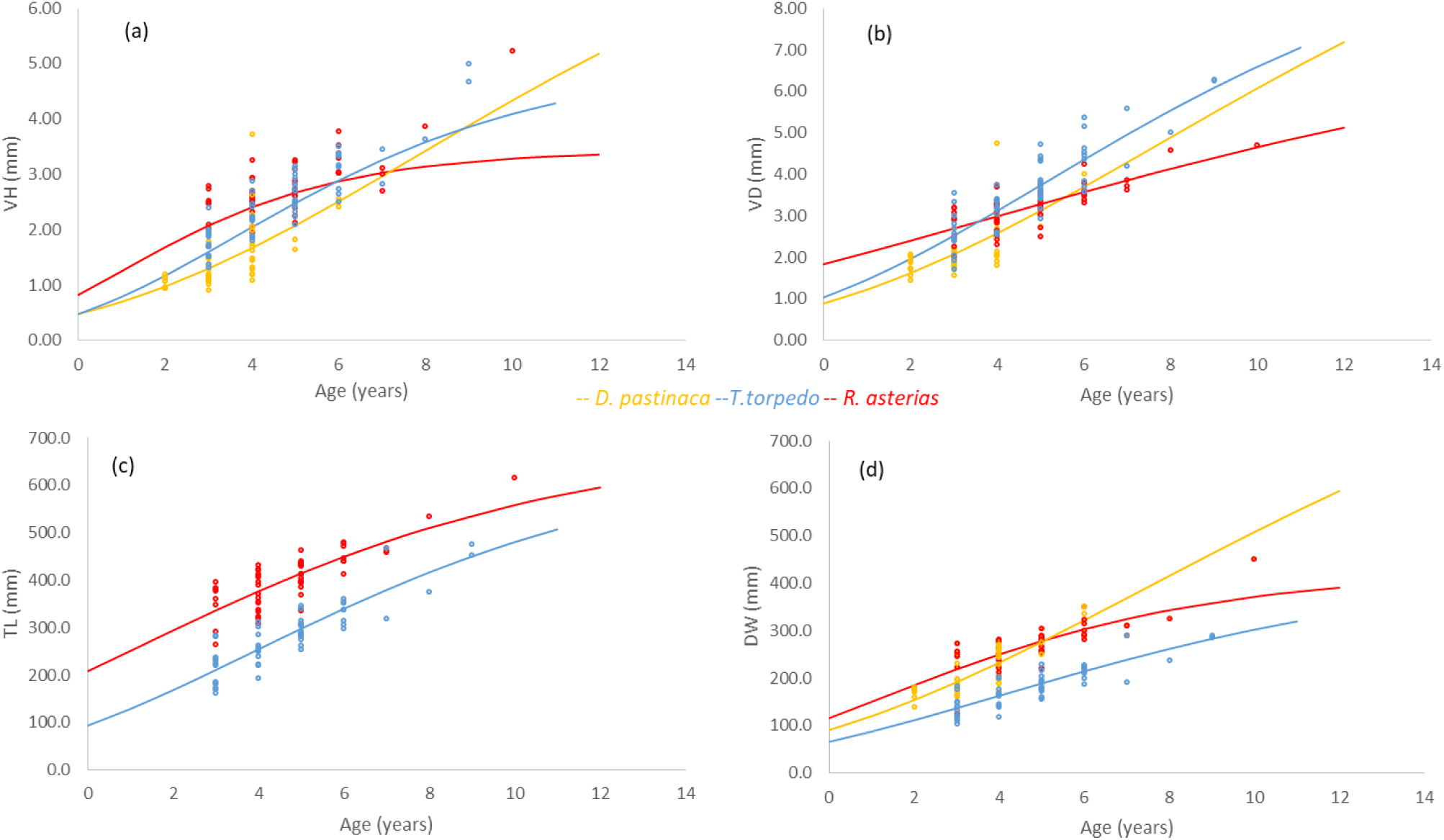
**a, b, c** Gompertz models based on vertebral (a) height (VH) and (b) diameter (VD), (c) total length (TL) and (d) disk width (DW) as asymptotic lengths in *Dasyatis pastinaca*, *Raja asterias* and *Torpedo torpedo* from the central Tyrrhenian Sea

## DISCUSSION

The study of the demographic status and population trajectory of species under fishing pressure requires reliable age-determination techniques (Matta et al., 2017), preferably rapid and inexpensive, as well as being multi-specific compared to those available. On the one hand, most ageing techniques involve staining procedures with expensive reagents (e.g., alizarin red, hematoxylin, xylene, crystal violet, graphite microtopography, cobalt nitrate, ammonium sulfide, Safranin O, Alcian blue, copper lead, iron-based salts, and silver nitrate), the operation of which depends strictly on the species being analyzed (Calliet et al., 1981; Price and Pulos, 1983; Gallagher et al., 2006; Basusta et al., 2010, 2017; Calliet, 2015; Matta et al., 2017). This is particularly true in the vertebrae of batoid species, as their vertebrae are difficult to read without histology or staining of some sort in many species (James and Natanson, 2020). On the other hand, the vertebrae of coastal species, such as carcharhinids and lamnids, do not require any stain techniques and can be read directly (James and Natanson, 2020). The most advanced use spectroscopy, from near-infrared spectroscopy to X-ray techniques (Calliet, 2015; Matta et al., 2017; Jung et al., 2018; Rigby et al., 2018; Arrington et al., 2021).

The method here presented allowed rapid and reliable reading of the vertebral age rings in four coastal batoid elasmobranchs. The use of an inexpensive and handy apparatus such as the Dino led Lite enhanced the action of multiple beams of led lights striking the specimen orthogonally. This enhanced the contrast of growth increments on vertebral sections simply immersed in glycerol as the only reagent. Another equally important achievement is the effectiveness of the statistical method developed to calculate age and growth parameters on size-selective and multi-specific fishery samples. Overall, the use of this method can be a great advantage, both in terms of cost-effectiveness and time taken to read age rings in different species, and in terms of feasibility in experimental field settings, above all to provide comparable demographic information.

Verification of the method came from the consistency of the data we found at three different levels.

First, the good general agreement among observers demonstrated the internal consistency of age readings within and between them, i.e., the measurements can be considered repeatable (Beamish and Fournier, 1981). However, different errors have occurred among species, generally concentrated in large individuals. This is expected in aging studies because of error propagation with increasing age (Calliet, 2015) and morphological differences between the vertebrae of elasmobranch species (Tortonese,1968). The latter is the case of *D. pastinaca*, which showed the greatest error in age counting on its poorly calcified vertebrae, as well as more acute in shape than the other species, as indicated by the VD / VH ratio. Like the deep-coned vertebrae of deep-sea shark species (Gennari and Scacco, 2007), more concave, poorly calcified vertebrae are more difficult to read than the less acute vertebrae (Claeson and Hilger, 2011; Smith et al., 2013; Pears et al., 2020; Matta et al., 2017; McMillan et al., 2017) due to the concealing effect of growth bands and the folding of the outer vertebral margin (Calliet, 2015). The latter, which can occur also in batoid species, can add a significant age-count bias in the oldest specimens such that histological staining preparations are required in these samples (James and Natanson, 2020).

Second, we found that as body size, and therefore age, increased vertebral size (height and diameter) increased in all species, as recommended for data verification in aging studies (Goldman, 2004). On the one hand, high age growth rates did not correspond to high vertebral growth rate with increasing body size in the species studied. Indeed, the starry skate showed an intermediate to low rate of increase in vertebral size with an increase in body size compared to the common torpedo. This may indicate different allometric growth of body parts between different species (James and Natanson, 2020). For example, skates have a wider “non-vertebral” portion of the body than torpedoes as the former have a developed rostrum, which is absent in the second group (Tortonese, 1956). *Dasyatis pastinaca*, on the other hand, showed very slow growth, both in body size with increasing age, and in the height of the vertebrae with increasing body size. In this species, the growth rate of all body parts is severely constrained by the low general metabolic rate observed in the longer-lived and gestation high-investment species (Dulvy and Reynolds, 1997; Cortés, 1998; 2002; Augustine et al., 2022). It is worth noting the invariant trend of the vertebral diameter-height ratio, which highlighted a general isometry in the vertebral growth of the species studied as their body size increased. Isometric vertebral growth may be a common trait for batoid species (Tortonese, 1956), due to their typical dorso-ventrally depressed body shape with pectoral fins enlarged to varying degrees (Claeson and Hilger, 2011; James and Natanson, 2020). Despite some slight differences found between species (in the case of the common stingray, as discussed above), the incomplete sample size range (juveniles and subadults, with few adult individuals) prevented stronger evidence to support this contention.

The third support to the proposed method comes from the reliability of the growth parameters that we estimated through the VBGE and Gompertz models applied, compared with information available in literature, although the samples suffered the consequences of a size-selective capture (Parma and Deriso, 1990). Indeed, they were not completely representative of the entire size range expected for the species considered, in addition to a limited and unbalanced sample size.

In this study, we addressed this problem by coupling the iteration of one of the simpler methods, the Ford-Walford method (1933, 1946), with a bootstrap routine applied to both vertebral and body measurements.

On the one hand, the iteration of the Ford-Walford method has helped to increase the accuracy of the asymptotic length estimation by providing a mean value with error. Indeed, while the Ford-Walford method suits problematic datasets (limited number of observations and lack of age groups), it provides estimates that are crude and serially correlated and depend on the sample size of the age groups (Ricker, 1975; Jones, 2000). On the other hand, bootstrapping helped account for all species-specific variability within the considered age classes (Ono et al., 2015; Goodyear, 2019), in particular, for those with small numbers of individuals. Finally, the asymptotic models with length constraints worked for the interdependence of the estimate of the growth parameters, as recommended by Ogle (2017), but not for the simultaneity of the latter. In the case under examination, the simultaneous estimates of asymptotic lengths, k and t_0_ would have suffered from a too wide error range due both to the low representativeness of the most numerous age groups (Tiralongo et al., 2018a, b), and the lack of the youngest individuals (0+ and 1 classes), i.e. a suboptimal situation for good adaptation (Jenkins and Quintana-Ascencio, 2020). We overpassed this problem by using the same samples first to calculate a set of mean asymptotic lengths and then to estimate k, kg, t_0_, cg through the asymptotic length-constrained VBGE and Gompertz models fitting the corresponding data. Estimation of growth parameters based on a set of mean asymptotic lengths between two model types enabled cross-variable and cross-model comparisons. Comparisons between the body variables and vertebral measures illustrated the former were better than the latter at estimating growth parameters with increasing age. In fact, the values of the rate of increment in vertebral dimension with increasing size not always had the same rank across species when considering the rate of increment in body measures with increasing age, as demonstrated in this study. Additionally, model comparisons confirmed the Gompertz as the best function for batoid species (Muller et al., 1995; Liu et al., 2021). A notable exception was *T. torpedo*, which showed good adaptations despite the model and/or variable used. This suggested that different measures and logistic functions can be used for this species for age and growth studies.

The resultant estimated growth rates k, as well as the asymptotic lengths, were generally in line with data available in the literature, and this is true for all species studied. For example, age data for *D. pastinaca* showed growth rate and mean asymptotic DW very close to data from Turkish (Ismen, 2003) and Greek (Yigin and Ismen, 2012) waters or higher and lower, respectively, than to the coastal waters of the Egyptian Mediterranean (Hamma, 1988). Large individuals are rare (Capapè et al., 2008) and Seret (2003) reported for this species a maximum disc width of 150 cm, about 50 cm greater than the asymptotic DW calculated in this study. The calculated age-growth rate for *T. torpedo* was consistent to other results on age-weight grow rate of the species from Spanish Mediterranean waters (Jaramillo-Londono et al., 2019) or higher than an outdated study from the Gulf of Tunis (Pauly, 1978). The species is reported at the maximum size of 40 cm TL in the southern Mediterranean (Abdel-Aziz, 1994), which is about 4 cm smaller than the one estimated here asymptotic TL. Age data for *R. asterias* showed that the actual sample grows almost as fast as a sample from a more northern area close to where it is today (Bono et al., 2005). On the other hand, the species appears to increase in weight and size in the northwestern Mediterranean, although the growth rate with age is not provided (Barria et al., 2015). Estimation of age growth parameters required caution for *T. marmorata*, due to limited sample size and sex composition (males only). In fact, while the asymptotic estimates appeared particularly lower in the marbled electric ray compared to the common torpedo, it is known that the former grows larger than the latter. The marbled electric ray grows up to 60 cm TL in the southern Mediterranean (Abdel-Aziz, 1994; Last et al., 2016) or even up to 100 cm as the maximum recorded TL (Møller, 1995). In fact, it exhibits a lower age and growth rate in Sardinian waters (Bellodi et al., 2021) than what was observed for the congener in this study. Even though the bootstrap routine used in present experiment functioned on difficult samples, the case of the marlbed electric ray indicated sample size cannot be too small and/or unbalanced for obtaining reliable age and growth estimates by using this method.

Although data provided in this work are not validated (Natanson et al., 2018), a condition that is achieved quite rarely (Calliet, 2015), age and growth data are comparable across species inhabiting the same area. In fact, all estimates were obtained using both a single band pair enhancing method and statistical routine on samples obtained from a local size selective fishery. This greatly reduced the biases connected with comparing samples with differences in statistical treatment and/or laboratory procedures of the aging methods used for different species. For instance, present data highlighted how the species-specific differences in the life history traits of the species studied can influence their age and growth rates. Indeed, the oviparous and more fecund starry skate showed the highest age and growth rate compared to the aplacental viviparous common torpedo, and, above all, to the common stingrays, a large sized species that evolved even intrauterine trophic connections with its embryos (Dulvy and Reynolds, 1997; Cortés, 1998; 2002; Augustine et al., 2022). Comparability of the data is equally important when studying populations of elasmobranch species distributed in different areas. In this case, intraspecific variation of the age and growth parameters from comparable data can inform on differences between factors that actually influence the species growth rate, such as biogeographic and bioecological conditions (trophic conditions, endemism, home range and gene flow, depending on the species) and in the fishing pressure exerted on populations in different areas (Kotas et al., 2011).

The impact of small-scale coastal fisheries on elasmobranch-dependent coastal species is a critical issue on a global scale (Heupel et al., 2007; Kinney and Simpfendorfer, 2009; Ward-Paige et al., 2014; Roff et al., 2018; Pacoureau et al., 2021), especially where the elasmobranch fauna is rich and composed of large endangered species (Temple et al., 2018). In fact, the IUCN red lists highlight how conservation measures are urgently needed for all dependent coastal species (IUCN 2021), in addition to obtaining local demographic assessments of populations under fishing pressure. In this context, studies on aging are essential to distinguish how the threat of fisheries affects populations, together with comparable demographic information that is critical for their conservation (Calliet, 2015). This information may be useful for addressing conservation measures on those age groups that are most affected by fishing within populations of elasmobranchs under fishing pressure (Matta et al., 2017). For example, restrictions on fishing in too shallow waters could help limiting the bycatch of the two-three years old juveniles and subadults that were the most represented individuals in the elasmobranch bycatch of the study area, as well as in similar areas, which are important feeding, breeding and/or nursery areas for many coastal elasmobranchs (Ward-Paige et al., 2014; Roff et al., 2018). As in this study, providing comparable demographic information for these species in an easily replicable method will help fill that information gap and help conserve them.

## Author Contributions

Conceptualization: [Umberto Scacco]; Methodology: [Umberto Scacco] and [Fabiana Zanardi]; Formal analysis and investigation: [Umberto Scacco] and [Silvio Kroha]; Writing - original draft preparation: [Umberto Scacco]; Writing - review and editing: [Umberto Scacco], [Francesco Tiralongo], [Emanuele Mancini] and [Silvio Kroha]; Funding acquisition: [Giuseppe Nascetti]; Resources: [Giuseppe Nascetti], [Umberto Scacco] and [Fabiana Zanardi]; Supervision: [Umberto Scacco].

## Data Availability

The datasets generated during and/or analysed during the current study are available from the corresponding author on reasonable request.

## Ethical approval

This study implemented a responsible collecting of specimens sampled from the ordinary by-catch of the small-scale local fishery. Collected samples were always individuals that died at the catch and/or during on board disentangling operations. We specifically instructed and encouraged fishermen to facilitate the release at sea of the alive individuals being caught, when in safe conditions due to the harmfulness of three out the four species under study. Collecting was in full accordance with Italian laws and regulations in this context and conducted under the responsibility of University of Tuscia, within a collaboration project between the University and the fishery coop of Fiora’s Port Canal.

## Acknowledgements

We are grateful to the anonymous referees who reviewed the manuscript and to all fishermen of Fiora’s Port Canal who collaborated with this study. We are grateful to A. Annunziatellis who provided us with the map.

