## Supplemental Figure 1 for "AGE READINGS AND ASSESSMENT IN COASTAL BATOID ELASMOBRANCHS FROM SMALL-SCALE SIZE-SELECTIVE FISHERY: THE IMPORTANCE OF DATA COMPARABILITY IN MULTISPECIFIC ASSEMBLAGES"

**DINO-LITE EDGE 3.0 (DIGITAL MICROSCOPE)**

Equipped with a high-quality 1.3- to 5-megapixel sensor with crystal-clear image quality and natural color reproduction, magnification up to 220 times. It comes with DinoCapture 2.0 software for Windows and DinoXcope for macOS, for images calibration and measurements. Built-in adjustable polarizer reduces glare and reflection on shiny objects.

- Aluminum stand
- - 8 white LEDs
- - Polarizing filter
- - USB
- - IR cut filter > 650nm
- - Functions for measurements

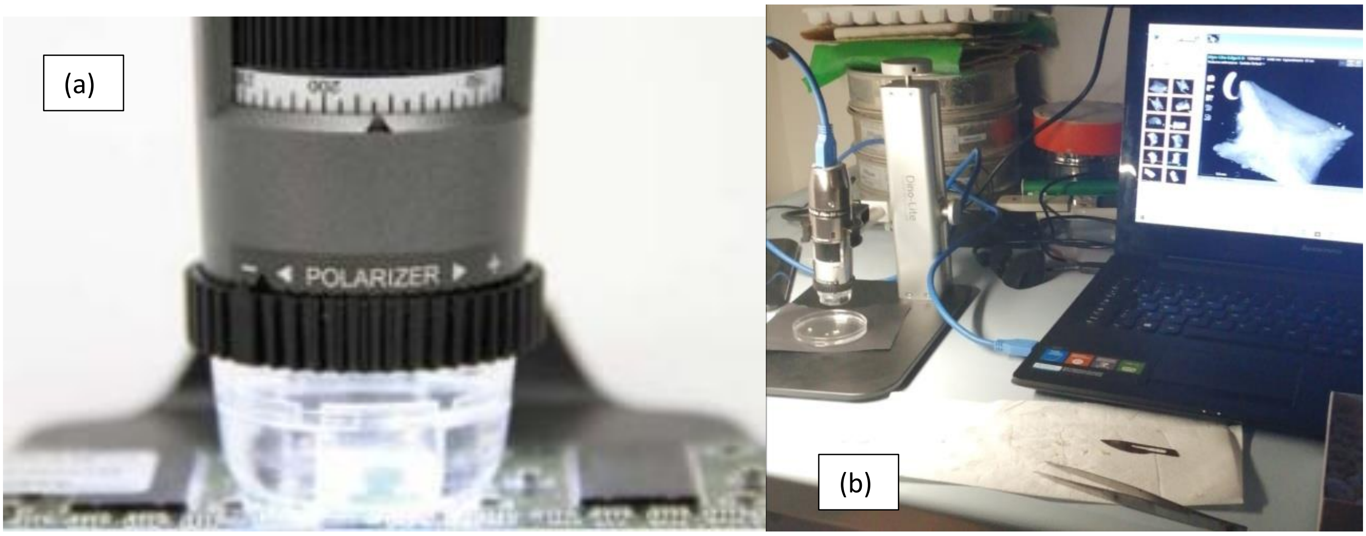


**Online information 1 a, b** Particulars of the used Dino Lite 3.0 used to observe the sagittal vertebral sections of four batoids species from the central Tyrrhenian sea, and to assess their age based on counts of growth bands on vertebral internal surface (a): details of the polarizer and magnification swift (b) detail of the stereomicroscope at work connected to a personal computer displaying an image of the vertebra under observation.
