## Supplemental Table 1 for "AGE READINGS AND ASSESSMENT IN COASTAL BATOID ELASMOBRANCHS FROM SMALL-SCALE SIZE-SELECTIVE FISHERY: THE IMPORTANCE OF DATA COMPARABILITY IN MULTISPECIFIC ASSEMBLAGES"

| Species | RP | *RA* | *TM* | *TT* |
|---|---|---|---|---|
| VD | ***a*** | $8.0*10^{-3}$ | $3.0*10^{-3}$ | $1.4*10^{-2}$ |
| | *Std. err* | $5.1*10^{-4}$ | $3.4*10^{-3}$ | $5.7*10^{-4}$ |
| | *t (n-2)* | 15.7 | 0.9 | 24.7 |
| | *p* | ***(a*) | ns | ***(a*) |
| | ***b*** | $2.0*10^{-6}$ | 2.79 | -0.40 |
| | *Std. err* | $2.1*10^{-1}$ | 0.96 | 0.16 |
| | *t (n-2)* | $1.0*10^{-4}$ | 2.9 | 2.4 |
| | *p* | ns | *** | * |
| | $R^2$ | 0.83 | $5.6*10^{-2}$ | 0.92 |
| VH | ***a*** | $7.7 *10^{-3}$ | $4.5*10^{-3}$ | $1.0*10^{-2}$ |
| | *Std. err* | $6.7 *10^{-4}$ | $3.6*10^{-3}$ | $4.5 *10^{-4}$ |
| | *t (n-2)* | 11.3 | 1.3 | 22.0 |
| | *p* | ***(b*) | ns | ***(b*) |
| | ***b*** | -0.33 | 1.45 | -0.32 |
| | *Std. err* | 0.29 | 1.0 | 0.13 |
| | *t (n-2)* | -1.2 | 1.4 | -2.4 |
| | *p* | ns | ns | * |
| | $R^2$ | 0.71 | 0.11 | 0.90 |
| VD/VH | **a** | $-4.4*10^{-5}$ | $-1.0*10^{-3}$ | $-1.0*10^{-5}$ |
| | Std. err | $3.1*10^{-4}$ | $1.5*10^{-3}$ | $2.6*10^{-4}$ |
| | t (n-2) | $-1.4*10^{-1}$ | $6.6*10^{-1}$ | $-4.0*10^{-2}$ |
| | p | ns | ns | ns |
| | **b** | 1.18 | 1.65 | 1.42 |
| | Std. err | 0.13 | 0.43 | $7.7*10^{-2}$ |
| | t (n-2) | 9.3 | 3.8 | 18.4 |
| | p | *** | ** | *** |
| | $R^2$ | $3.9*10^{-4}$ | $3.3*10^{-2}$ | $3.0*10^{-5}$ |

**Online information 2** Regression parameters (RP, a: slope, b: intercept, $R^2$: explained variance, Std. err: standard error, *t (n-2)*: t statistic for n-2 degrees of freedom with n as sample size, and p: level of significance) of linear relationships between vertebral dimensions and increasing fish size as total length (mm) of three coastal batoid species (RA: *Raja asterias*, TM: *Torpedo marmorata* and TT: *Torpedo torpedo*) sampled in the central Tyrrhenian Sea. VD, VH and VD/VH are vertebral diameter, height and their ratio, respectively. Same letter in brackets indicates significantly different regression slope between species with asterisks denoting p-level of significance of difference in slope in pairwise comparisons between species. ns stands for not significant differences. * $p < 0.05$, ** $p < 0.01$
