## Supplemental Figure 3 for "AGE READINGS AND ASSESSMENT IN COASTAL BATOID ELASMOBRANCHS FROM SMALL-SCALE SIZE-SELECTIVE FISHERY: THE IMPORTANCE OF DATA COMPARABILITY IN MULTISPECIFIC ASSEMBLAGES"

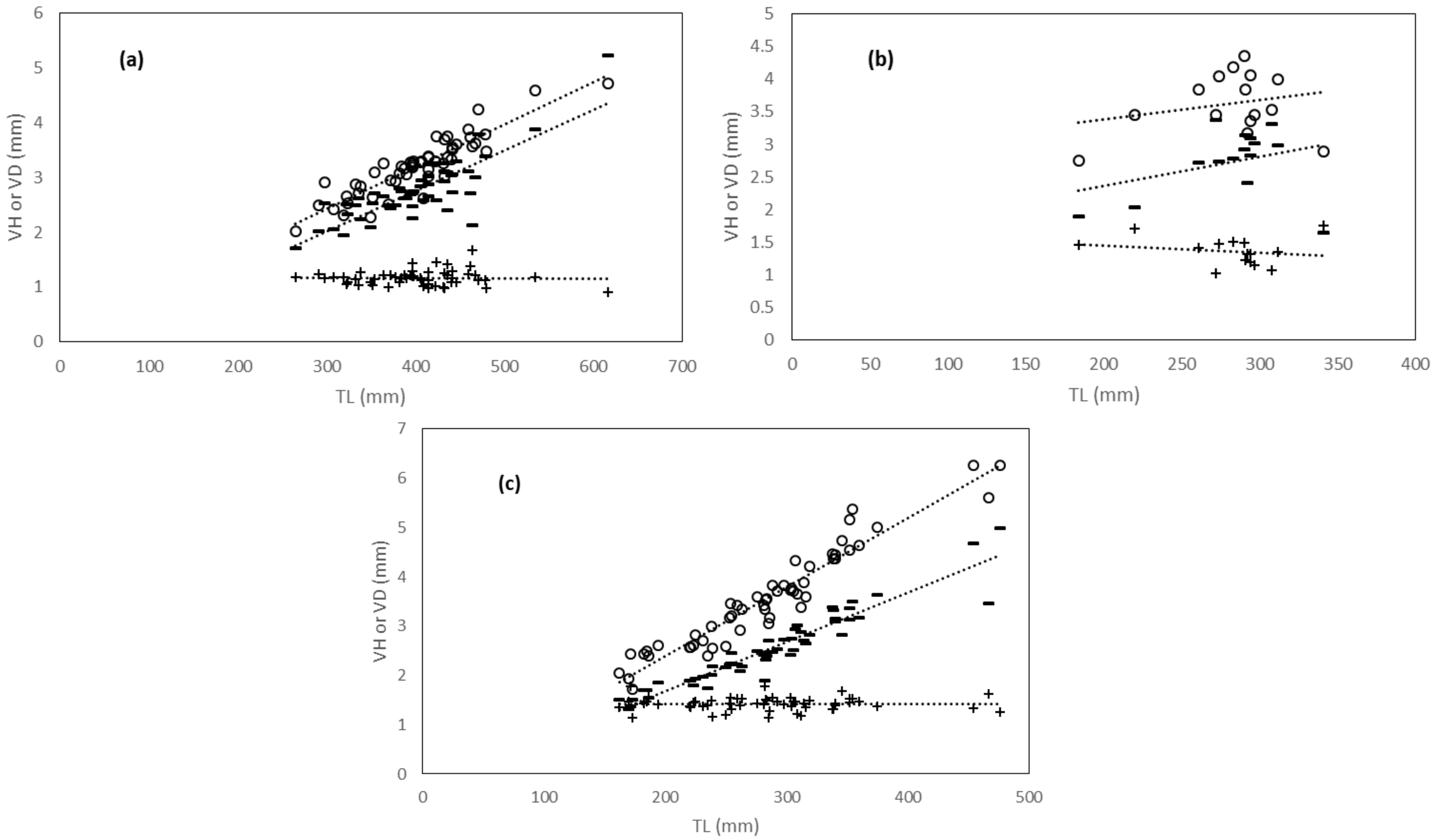


**Online information 4 a, b, c** Linear relationships between total length (TL) and vertebral measures (VD vertebral diameter, empty circles; VH vertebral height, dark lines; VD/VH dimeter/height ratio, dark crosses) in three coastal batoid elasmobranch species form the central Tyrrhenian sea. (a): *Raja asterias*; (b): *Torpedo marmorata*; (c): *Torpedo torpedo*
