## Supplemental Figure 4 for "AGE READINGS AND ASSESSMENT IN COASTAL BATOID ELASMOBRANCHS FROM SMALL-SCALE SIZE-SELECTIVE FISHERY: THE IMPORTANCE OF DATA COMPARABILITY IN MULTISPECIFIC ASSEMBLAGES"

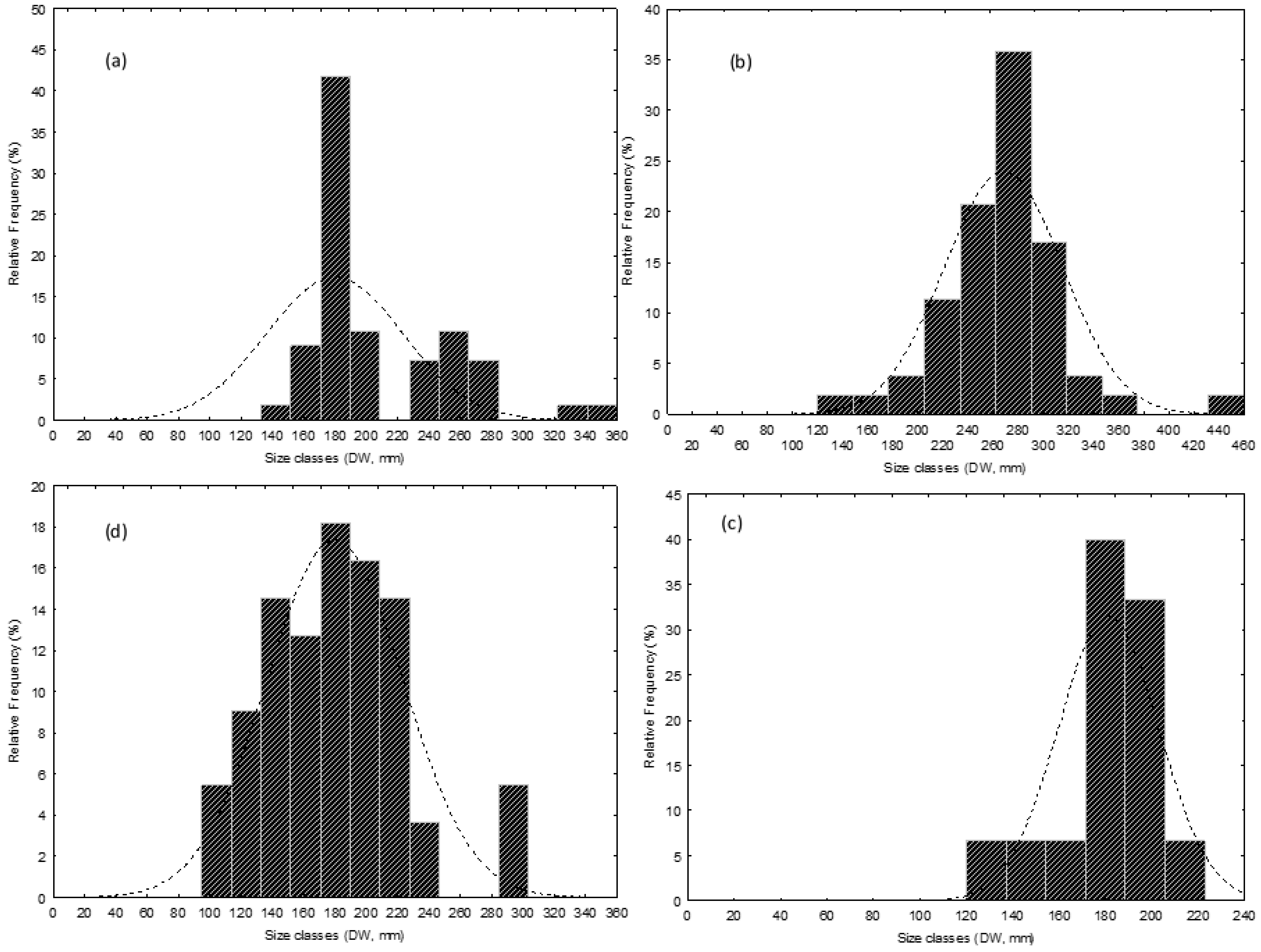

(a)
Relative Frequency (%)
0 5 10 15 20 25 30 35 40 45 50
0 20 40 60 80 100 120 140 160 180 200 220 240 260 280 300 320 340 360
Size classes (DW, mm)
(b)
Relative Frequency (%)
0 5 10 15 20 25 30 35 40
0 40 80 120 160 200 240 280 320 360 400 440
20 60 100 140 180 220 260 300 340 380 420 460
Size classes (DW, mm)
(d)
Relative Frequency (%)
0 2 4 6 8 10 12 14 16 18 20
0 20 40 60 80 100 120 140 160 180 200 220 240 260 280 300 320 340 360
Size classes (DW, mm)
(c)
Relative Frequency (%)
0 5 10 15 20 25 30 35 40 45
0 20 40 60 80 100 120 140 160 180 200 220 240
Size classes (DW, mm)

**Online information 5 a, b, c, d** Size-frequency distributions of four coastal batoid elasmobranch species from the bycatch of size-selective small scale fishery form the central Tyrrhenian sea. (a): *Dasyatis pastinaca*; (b): *Raja asterias*; (c): *Torpedo marmorata*; (d): *Torpedo torpedo*. DW is disk width in mm.
