## Supplemental Figure 5 for "AGE READINGS AND ASSESSMENT IN COASTAL BATOID ELASMOBRANCHS FROM SMALL-SCALE SIZE-SELECTIVE FISHERY: THE IMPORTANCE OF DATA COMPARABILITY IN MULTISPECIFIC ASSEMBLAGES"

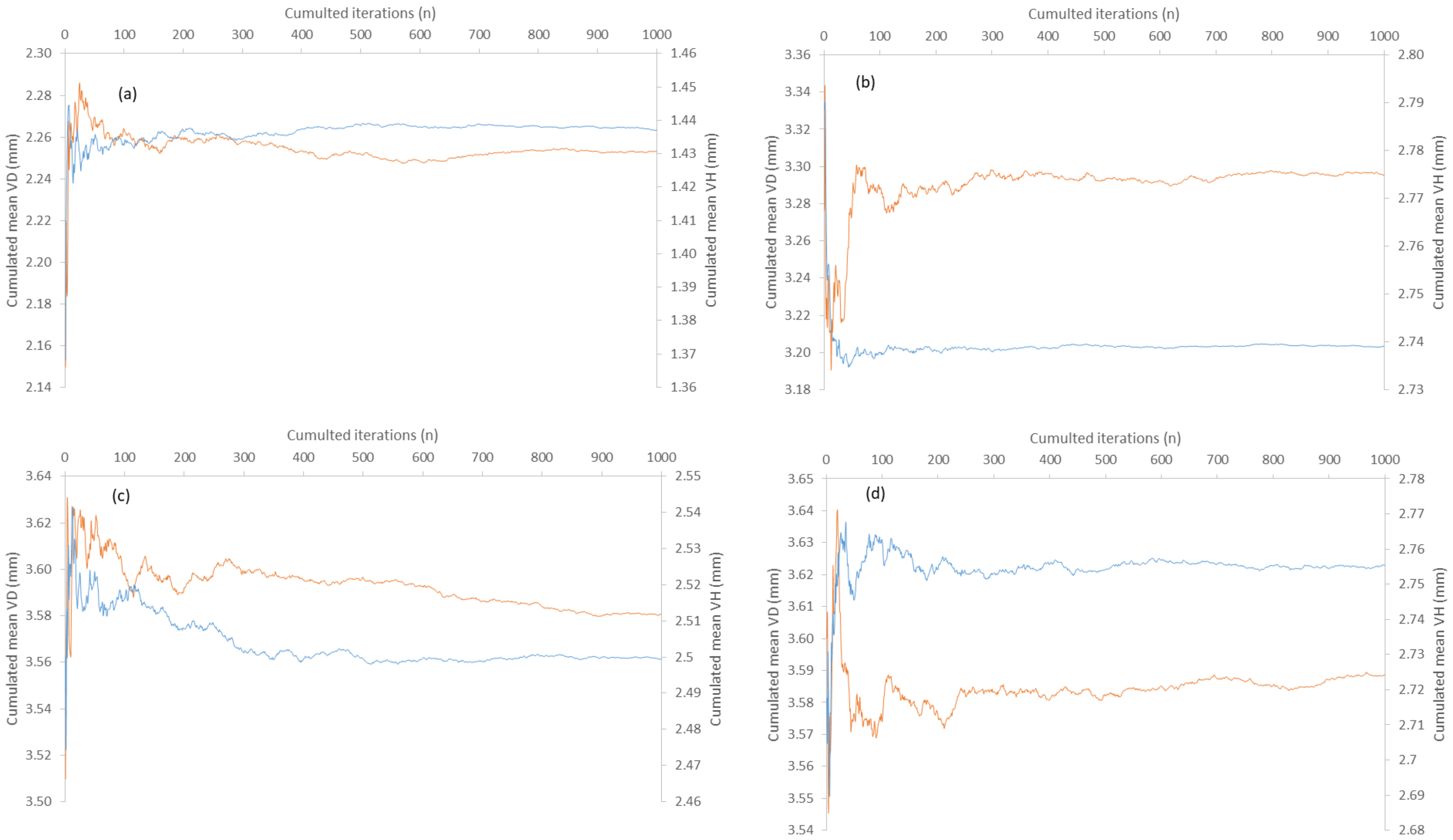


**Online information 5 a, b, c, d** Number of cumulated iterations needed for stabilizing cumulated sample mean of vertebral height (VH) and diameter (VD) in the bootstrapping procedure applied to: (a) *Dasyatis pastinaca*, (b) *Raja asterias*, (c) *Torpedo torpedo* and (d) *Torpedo marmorata* sampled in coastal waters of the central Tyrrhenian Sea.

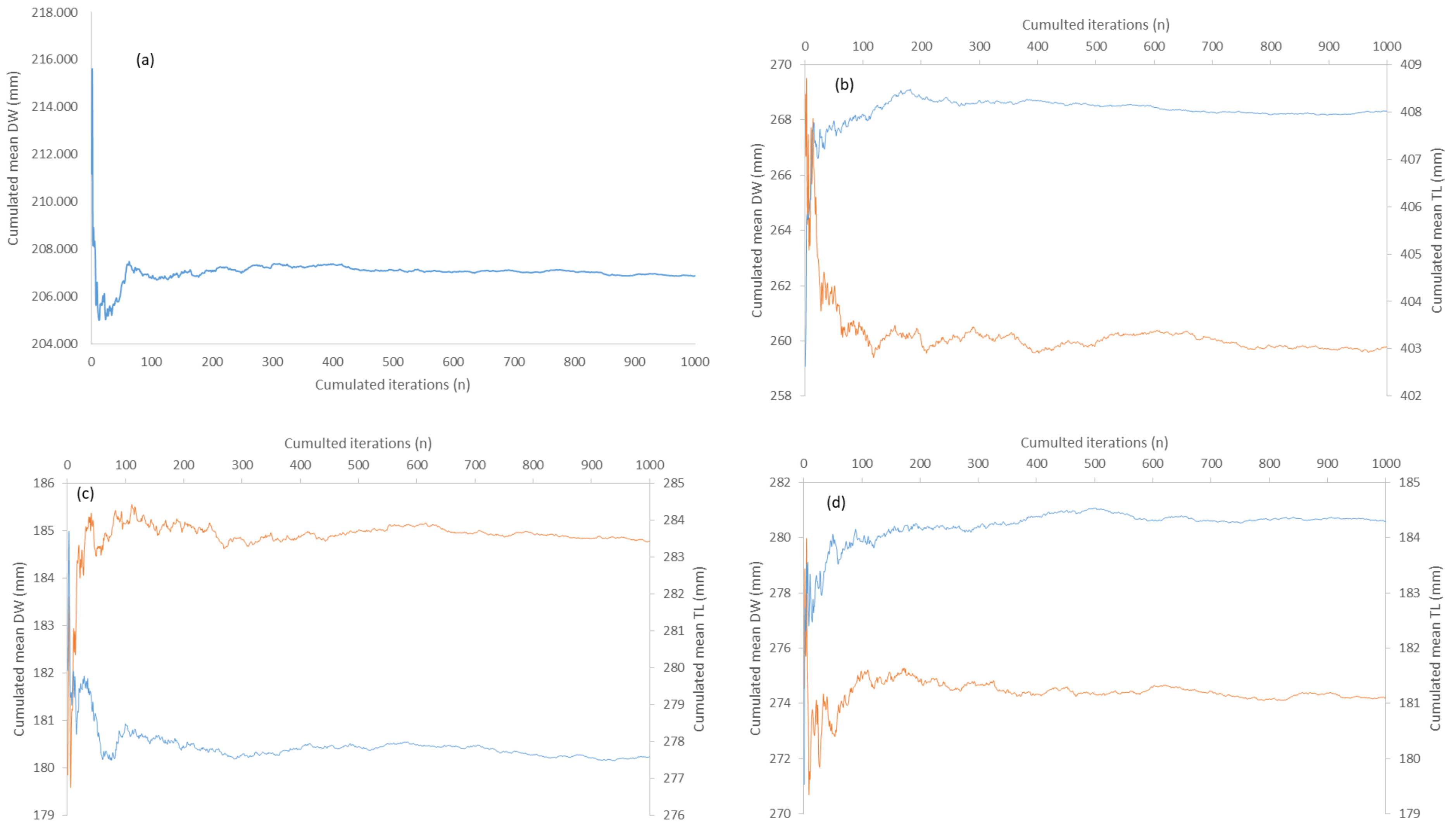

(a)
Cumulated mean DW (mm)
Cumulated iterations (n)
218.000
216.000
214.000
212.000
210.000
208.000
206.000
204.000
0
100
200
300
400
500
600
700
800
900
1000
(b)
Cumulted iterations (n)
Cumulated mean DW (mm)
Cumulated mean TL (mm)
270
268
266
264
262
260
258
409
408
407
406
405
404
403
402
(c)
Cumulted iterations (n)
Cumulated mean DW (mm)
Cumulated mean TL (mm)
186
185
184
183
182
181
180
179
285
284
283
282
281
280
279
278
277
276
(d)
Cumulted iterations (n)
Cumulated mean DW (mm)
Cumulated mean TL (mm)
282
280
278
276
274
272
270
185
184
183
182
181
180
179

**Online information 5 bis a, b, c, d** Number of cumulated iterations needed for stabilizing cumulated sample mean of disk width (DW) and total length (TL) in the bootstrapping procedure applied to: (a) *Dasyatis pastinaca*, (b) *Raja asterias*, (c) *Torpedo torpedo* and (d*) Torpedo marmorata* sampled in coastal waters of the central Tyrrhenian Sea.
